# Information, Movement and Adaptation in Human Vision

**DOI:** 10.1101/2025.09.05.674315

**Authors:** Alexander J.H. Houston, David H. Brainard, Hannah E. Smithson, Daniel J. Read

## Abstract

Our eyes are never still. Even when fixating, they exhibit small, jittery motions. While it has long been argued that these fixational eye movements (FEMs) aid the acquisition of visual information, a complete theoretical description of their impact on the information available in the early visual system has been lacking. Here we build FEMs into a minimal theoretical model of the early visual response, including the critical process of adaptation (a fading response to a fixed image). We establish the effect of FEMs on the mutual information between a visual stimulus and this response. Our approach identifies the key dimensionless parameters that characterise the effect of fixational eye movements, and reveals the regimes in which this effect can be beneficial, detrimental or negligible. Taking parameter values appropriate for human vision, we show that the spatio-temporal couplings induced by fixational eye movements can explain the qualitative features of the human contrast sensitivity function as well as classic experiments on temporal integration. To our knowledge, the consideration of fixational eye movements in this latter context is novel, and our results suggest the need for future experiments to determine the mechanisms by which spatial and temporal responses are coupled in the human visual system.

## I. INTRODUCTION

Human vision is a dynamic process, and our eyes are in constant motion [1]. At the largest scales, as we look around a scene, jump-like movements of the eye called saccades locate the images of targets of interest on the high-resolution fovea in the centre of the retina [2]. In between saccades we fixate on a target, and yet our eyes still move. These fixational eye movements (FEMs) comprise occasional small jumps of order 10 arcmin known as microsaccades, but for the majority of the time the eye undergoes a seemingly random, diffusion-like motion referred to as drift [1, 3, 4]. That the eye performs fixational movements has been known for over 250 years, thanks to the observations of Jurin [5], Robert Darwin [6] and Helmholtz [7], who made reference to the ‘wandering of the gaze’. As to why these movements occur, a pessimistic view is that FEMs are merely involuntary motions resulting from imperfect motor control of the eye which simply blur or degrade an image [8]. However, this view seems unlikely to provide a full explanation, given the many aspects of vision which have evolved to achieve optimal performance given some physical constraint [9, 10], such as single photon detection [11–13], the diffraction-limited construction of insect eyes [14]; discussed by Feynman [15] and McLeish [16], and the use of Cartesian ovals in Trilobite eyes to correct abberation [17]. Indeed, the idea that FEMs may be beneficial dates back to 1899 [18] and there has been growing empirical evidence that they can improve visual acuity [19–21]. Further, it has been suggested that the magnitude of drift can be tuned [22–24], such that the view of FEMs has developed from that of an aimless ‘wandering eye’ to a more purposeful, controlled ‘active vision’ [25, 26].

How, then, might FEMs be beneficial, how should this benefit be optimised, and in what fashion does it depend on the features of the visual stimulus? The visual system is designed to detect changes. So much so that, were the environment completely static and our eyes completely still, we would not see anything at all. This visual adaptation, which functions as a band-pass temporal response, has been well-established [27, 28] and opens the door for FEMs to beneficially refresh the image by moving the stimulus over the retina. However, the effect of FEMs is not uniform for all spatial frequencies with an image. Rather they cause the spatial structure of a static image to be converted into spatio-temporal structure in a manner that depends on the spatial frequency [1, 29]. Accordingly, it has been hypothesised that FEMs serve not simply to overcome adaptation by refreshing the image, but to encode visual information within a spatio-temporal structure to which subsequent processing stages are most sensitive [18, 30–34]. Some early psychophysical experiments [35] did not support this view, however, and there remains debate as to the degree to which human visual performance can be understood without considering FEMs [36] versus only with the incorporation their effects [26].

To address the issue of how FEMs might be expected to affect visual performance, here we develop an analytically tractable model of the effect of drift on the information about a stimulus available in the early visual system. The model exposes basic principles of how stimulus properties (spatial frequency, temporal frequency, duration), spatial and temporal filtering in the visual pathways, and properties of fixational eye movements (rate of drift) interact to shape the limits of visual performance. Related to this, the model derives key dimensionless quantities that capture aspects of these interactions and allows quantitative prediction of how visual performance depends on both the stimulus and key components of the neural processing chain. One key implication of the model is that across a range of stimulus parameters (e.g. duration and spatial frequency), it is possible to find regimes in which FEMs enable increased performance and other regimes in which they are deleterious. The existence of both types of regime illustrates the richness of the interplay between FEMs and the visual information available for perceptual decisions, and clarifies why effects of both signs can potentially be observed in experimental data.

The thread which runs through this paper is that spatial variation in a stimulus combined with retinal motion produces a time-dependent visual signal. Although simple, we find this principle to have great expository power. We make it quantitative in Section II by constructing a minimal model for the early stages of the visual system, allowing theoretical investigation of the influence of ocular drift on information acquired about an external stimulus.

In Section III A we use the model to determine the optimal form of diffusive eye motion to gain information from a given spatial wave vector. Taking this further, we establish the combinations of stimulus presentation time and spatial frequency for which ocular drift is beneficial, detrimental or has negligible effect on sensitivity to individual spatial wave vectors. Although we do not attempt a detailed account of specific experiments on human contrast spatial sensitivity, by employing reasonable numerical estimates of properties of the human visual system we find that measured rates of fixational ocular drift are tuned so as to provide maximum benefit given the presence of optical blur - an indication that once again a biological process is operating near a limit imposed by physical principles.

In Section III B we incorporate temporal modulation of the stimulus. Here we find that the fact that eye movements convert spatial variations into temporal ones, in conjunction with temporal adaptation and blur, is able to account qualitatively for the features of the spatiotemporal contrast sensitivity function without the need to invoke a bandpass spatial filtering stage. This is not to imply that such stages do not exist nor that a quantitative account can be achieved without incorporating them, but rather to again illustrate the range of effects that can be understood when the interactions between FEMs and visual system filtering are jointly considered in a systematic framework.

We consider a second example in Section IV, the extent to which the experiments of Barlow [37] support the notion that the temporal adaptation of the human visual system is dependent on the scale of the stimulus, as has been understood previously. By using a theoretical model of detection [38], we again find that the modulation of information caused by ocular drift is sufficient to explain the observations, without incorporating additional layers of post-receptoral processing. A given form of eye motion will induce temporal signal variations at a frequency dependent on the spatial scale of the stimulus, and this results in an effective scale-dependent adaptation which allows the features of the experimental observations to be reproduced. As with our analysis of spatio-temporal constrast sensitivity, our point is not that the inferred effects are necessarily absent, but rather that caution is required before the inferences are secure. Our analysis in this case suggests the structure of future experiments that might be decisive.

We close with a brief Discussion.

## II. INFORMATION IN THE EARLY VISUAL SYSTEM

### A. Minimal model of the early visual response

Our aim in this paper is to understand the role that fixational eye movements play in determining the visual information available at the retina. Our approach is to employ a minimal model of the early visual system. We take the position that much of the impact of FEMs stems from their coupling of spatial and temporal properties, specifically the spatial structure of the stimulus and a band-pass temporal response. Therefore, while we incorporate a temporal adaptation filter, we neglect any additional aspects of post-retinal signal processing, although these could be incorporated into our model, as discussed in Section V. To further simplify the theoretical description, we treat the photoreceptor array as a homogeneous continuum of constant photoreceptor density, neglecting that photoreceptors are discrete objects and also the preferential response of L, M and S cones to long, medium or short wavelengths. We consider only contrast stimuli, to which we assume all photoreceptors have an identical response. While the band-pass temporal response does not occur in the photoreceptors, in the interest of concision, we shall refer to our minimal model of the response of the early visual system as the ‘photoreceptor response’.

The key components of our model of photoreceptor response are laid out in Figure 1. There are three ways in which time-dependence of the photoreceptor signal, *y*, can be induced: temporal modulation of the external stimulus, *x*, retinal motion, and adaptation. The first of these is described by an amplitude *A*(*t*^*′*^), with the example shown in Figure 1(b) corresponding to a stationary external stimulus presented with constant amplitude for a time *T*. Note that we take this modulation to be uniform across the stimulus, a common experimental protocol, such that the spatial and temporal variation separate and hence appear as the product *x*(**r**^*′*^)*A*(*t*^*′*^) in (1). As a convention we take |*A*(*t*^*′*^)| ≤ 1, with the overall stimulus amplitude incorporated into *x*(**r**^*′*^).

**FIG. 1.**
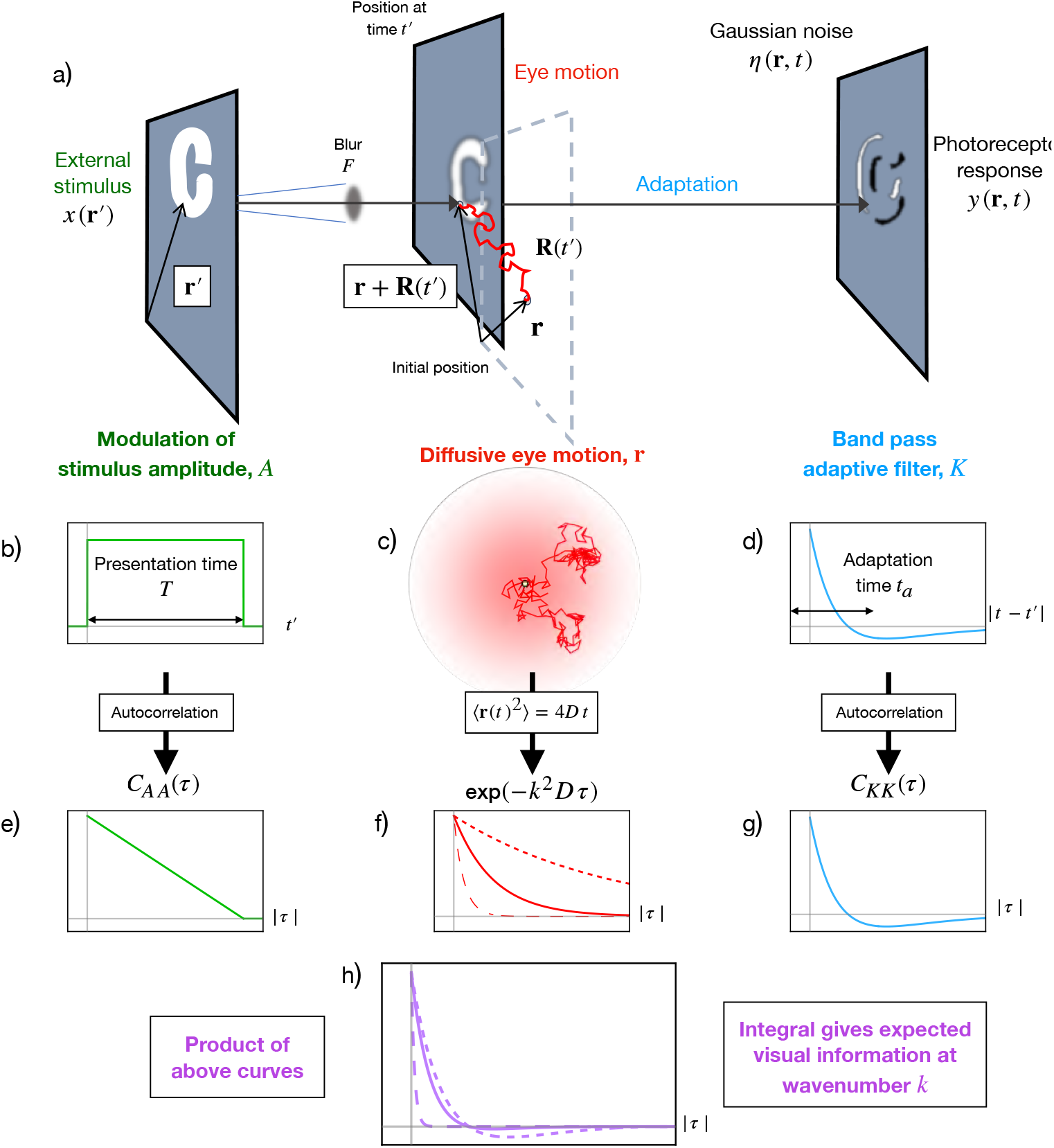
A schematic of our minimal model for the visual information acquired from a stimulus. a) An external stimulus is related to a photoreceptor response through optical blur, eye motion, adaptation and Gaussian noise. Setting aside blur for the moment, in our minimal model the central elements of the stimulus, eye motion and retinal physiology are captured by the stimulus amplitude, diffusive trajectories and a linear adaptive response, which are shown in b), c) and d) respectively. (Although we focus on diffusive eye motion in the paper, our model allows more general motion to be incorporated.) In particular, it follows from a calculation presented in the SI that the aspects of the stimulus, eye motion and adaptive response that are relevant for the acquisition of visual information are the functions shown in e), f) and g), where *C*_*AA*_ and *C*_*KK*_ denote the autocorrelations of the stimulus and adaptive filter and 4*Dt* is the mean squared displacement of the retina, with *D* the diffusion constant. The solid, small dashed and large dashed curves correspond to a medium, small and large value of *k*^2^*D*, where *k* is a spatial wavenumber of the stimulus. The product of the functions shown in e), f) and g) produces the curves shown in h), the integral of which gives the visual information acquired from the stimulus at wavenumber *k*, up to a given time, modulo additional effects due to blur as discussed in the main text. The different curves in h) result from the corresponding curves in f).

Secondly, retinal motion will convert spatial variation of the stimulus into temporal variation of the photoreceptor response. A photoreceptor initially located at **r** will, at some later time *t*^*′*^, have moved to a new position **r** + **R**(*t*^*′*^) via retinal motion encoded by **R**(*t*^*′*^). Due to the optical blur of the eye’s lens, and because of its finite size, a photoreceptor will collect photons originating from a distribution of positions within the stimulus. We represent this distribution by a blurring function *F* centred on the current photoreceptor position, leading to the convolution between the stimulus and blurring function shown in Eq. (1). In what follows we assume that the path **R**(*t*^*′*^) taken by the eye is ‘known’ to the neural processors, either via mechanical or electrical signals or by inference from a wider set of visual cues, and hence this motion is not itself a source of noise. The inference of motion is itself an interesting area of study [39, 40]. Here we simply assume that there is sufficient data available for this inference to be accurately made.

Lastly, the temporal response of photoreceptors (and early stage signal processing in the retina) is not instantaneous. These early components of the visual signaling pathway integrate an incoming signal over some past time to accumulate a response [38], and adapt to steady stimuli such that a constant stimulus will eventually produce a zero, or near zero response. This can be modelled by a linear temporal response function, or memory kernel, *K*(*t* −*t*^*′*^) [38]. The present photoreceptor response at time *t* corresponds to integration over the past signal from earlier times *t*^*′*^, weighted by the memory kernel, as can be seen below in Eq. (1). Typically the memory kernel is considered to be positive for small *t* −*t*^*′*^, to model the accumulation of input signal at short times, before becoming negative for larger *t* −*t*^*′*^ to model the adaptation process. For perfect adaptation, that is zero response to a constant signal in the long time limit, the integral of *K* should be zero. These features are apparent in the example shown in Figure 1(d), which takes *K* to be bi-exponential. We shall use this form of memory kernel throughout this work and will describe it more fully shortly.

The preceding elements, with the inclusion of additive noise, combine to give the following minimal model for the photoreceptor array response

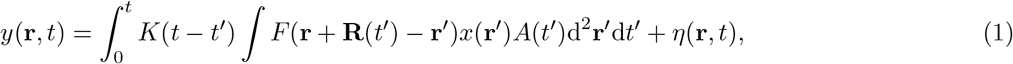

as has been used previously in the context of motion estimation [39]. We reiterate that (1) provides the response at time *t* of a photoreceptor initially located at **r**. We take the noise *η*(**r**, *t*) to be Gaussian white noise, i.e. uncorrelated between different points in space or time. One could also include noise on the stimulus *x*(**r**^*′*^), for example due to photon shot noise, which becomes a significant factor at low light levels [41–44], or noise in the early stages of photoreceptor excitations. Such noise would be passed through some of the temporal and spatial filtering represented by the functions *K* and *F*, resulting in coloured noise at the final signal. We neglect this for simplicity.

### B. Quantifying visual information

We are now in a position to consider the rate at which visual information can be gained about the external stimulus *x* from the photoreceptor response *y*. This process is hindered by noise, adaptation and optical blur. A central question, which our model allows us to address, is to what extent eye movements mitigate these factors. It is convenient to take the spatial Fourier transform of Eq. (1) and to consider the photoreceptor array response at wavevector **k** (i.e. with a spatial frequency |**k**|*/*2*π*). Denoting the Fourier transform by 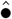, this gives 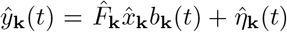, where 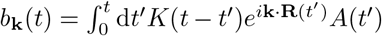. Maximising the information transmitted at wavevector **k** corresponds to making *b*_**k**_(*t*), in some sense, as large as possible. We have found that the appropriate measure of the magnitude of *b*_**k**_(*t*) is the quantity

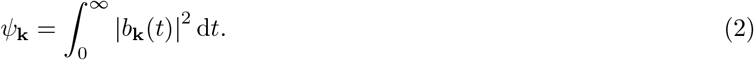

Making *ψ*_**k**_ large corresponds to making the signal at wavevector **k** large, in a time integrated sense. Once the other components of the visual system have been specified, the only remaining optimisation problem is how to maximise this quantity via eye movements. In the Section I of the Supplemental Material [45], we analyse multiple different ways of quantifying the process of learning about external stimulus *x* from the photoreceptor response *y*, and in each case demonstrate that *ψ*_**k**_ emerges as the correct quantity to consider. We show that maximising *ψ*_**k**_ minimises the mean square error when estimating the stimulus 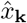 from the response 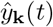. A robust and often-used measure is the mutual information: assuming a Gaussian prior distribution of the stimulus 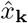, we show that maximising *ψ*_**k**_ gives maximal mutual information between stimulus *x* and photoreceptor response *y* at wavevector **k**. We finally show that this same quantity *ψ*_**k**_ emerges from an existing model of signal detection [38], that we will employ in Section IV. Note that in all cases that the information transmitted depends on the product 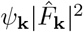, and in this way depends also on the optical blur. This simple dependence on the blur function arises because we assume *F* does not have a temporal component (i.e. that there is a factorisation into separate functions governing spatial (*F*) and temporal (*K*) response).

The value of *ψ*_**k**_ depends on the specific path **R**(*t*^*′*^) of the eye motion, i.e. it takes a different value for different paths. In the present work we focus on its average ⟨*ψ*_**k**_⟩ across multiple paths, reasoning that forms of eye motion that maximise the average are likely to be beneficial. Here and throughout we use ⟨•⟩ to denote an average over the ensemble of possible paths taken by the eye. Under the assumption that the eye movements are a statistically stationary process, meaning that there is no special reference time and so the statistical properties of the motion do not vary over time, we show in Section II A of the Supplemental Material [45] that the average value of *ψ*_**k**_ can be conveniently written as

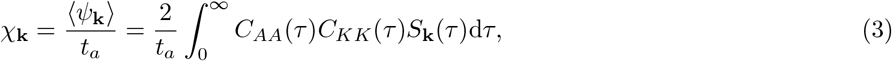

where *C*_*ff*_ denotes the auto-correlation of the function *f*,

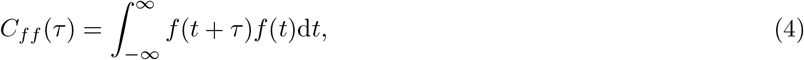

*S*_**k**_(*τ*) = ⟨exp [i**k** · (**R**(*t* + *τ*) ™ **R**(*t*))] ⟩ and *t*_*a*_ is an adaptation time that we define in the following paragraph. The three terms under the integral for *χ*_**k**_ in (3) capture the effect on information gain of the three sources of time-dependence in the photoreceptor response, these being temporal modulation of the stimulus, adaptation and eye movements, as discussed previously. Typical forms of these three terms are shown in Figure 1, and we now describe each in turn.

We begin with the contribution to the retinal information due to the memory kernel. Since, in our minimal model, the memory kernel captures the retinal physiology, it provides the relevant timescale by which to normalise all others. As mentioned in II A, throughout this work we take the memory kernel to be bi-exponential, as shown in in Figure 1(d). It is composed of an excitatory response on a short timescale *t*_s_ and an inhibatory one on a long timescale *t*_l_. From these we define two auxillary parameters: an ‘adaptation time’ 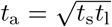 and a ‘shape parameter’ 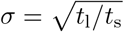. The adaptation time sets the characteristic timescale for the memory kernel and it is useful to use it to non-dimensionalise other times in the system, which we shall denote with a tilde, for example 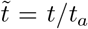. The shape parameter encodes the separation between short and long time responses of the temporal filter (so, *t*_s_ = *t*_a_*/σ* and *t*_l_ = *t*_a_*σ*). With this choice of parameters the memory kernel is rendered pleasingly symmetric

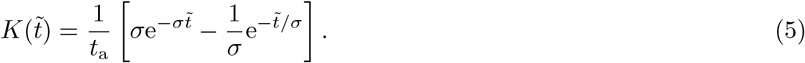

for 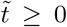, with 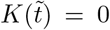 for 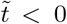. As presented in (5) the memory function exhibits perfect adaptation; the excitatory and inhibatory components are balanced such that at long times the response to a constant stimulus will vanish. Imperfect adaptation can be incorporated by giving the two components different weightings. We discuss the consequences of this in Section IV of the Supplemental Material [45]. For a bi-exponential memory kernel the autocorrelation, as calculated via (4), produces a scaled version of the original function

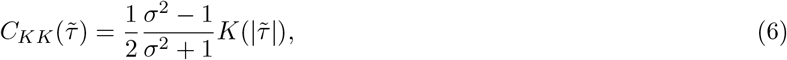

as illustrated in Figure 1(g). The prefactor of *C*_*KK*_ reflects that when *σ* = 1 the two timescales are equal and the memory kernel vanishes.

Some previous studies have used a higher order memory kernel as results from successive application of exponential filters [38, 46]. This can result in a markedly different memory kernel for which *K*(0) = 0. We note, however, that this difference is suppressed in the autocorrelation function, which remains qualitatively similar in profile to the bi-exponential in (6), as illustrated in Section III of the Supplemental Material [45] for a memory kernel of the form given by Watson [38]. In particular, the autocorrelation function in both cases is maximal at the origin and has a single minimum. Consequently, none of the results presented in this paper are qualitatively changed by using a higher order memory kernel.

The contribution of the stimulus temporal modulation to the retinal information is also captured via its autocorrelation function, again calculated via (4). For a constant amplitude stimulus presented for time *T* we take *A*(*t*) to be 1 when 0 ≤ *t* ≤ *T* and zero otherwise. For non-dimensional time 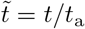 and presentation time 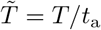, this gives 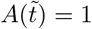 when 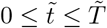 and zero otherwise, as shown in Figure 1(b). The resulting autocorrelation function is

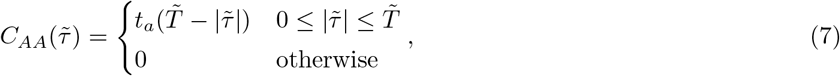

which has the decreasing triangular form shown in Figure 1(e). The autocorrelation function for sinusoidal stimulus modulation is shown in Section V of the Supplemental Material [45].

The impact of retinal motion on gaining visual information is captured by *S*_**k**_(*τ*) = ⟨e^i**k***·*(**R**(*t*+*τ*)*−***R**(*t*))^⟩. This path ‘structure factor’ involves summing the signal amplitudes due to different retinal positions, averaging over all paths that might occur and averaging over time *t*. We emphasise that this expression for *S*_**k**_ is valid for all forms of eye movement that are statistically stationary (i.e. whose statistical properties are independent of time). In the present work, following earlier studies [1], we treat the eye’s motion as diffusive (Figure 1(c)). This means that the probability distribution of motion vectors is zero-mean Gaussian, so that the path structure factor is specified wholly by the mean squared displacement (MSD) as 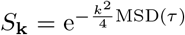. Diffusive motion is additionally constrained so that MSD is proportional to time, MSD(*τ*) = 4*Dτ*, with *D* the diffusion constant. Hence the expression used throughout this paper is

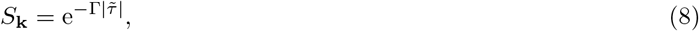

where we define the dimensionless parameter Γ = *k*^2^MSD(*t*_a_)*/*4 = *k*^2^*Dt*_a_, the square of the spatial frequency, made dimensionless by the diffusion constant and adaptation time. At fixed diffusion constant *D*, increasing Γ corresponds to increasing the square of the spatial frequency *k*; at fixed spatial frequency, increasing Γ corresponds to increasing the diffusion constant.

The logic behind the exponential decay, shown in Figure 1(f), is then clear. The dimensionless number Γ quantifies the typical magnitude of diffusive motion relative to the wavenumber *k* during the adaptation time *t*_a_, and hence determines the rapidity of the temporal variation of the photoreceptor signal induced by diffusive eye motion. The faster this variation, the smaller the amplitude that is produced upon averaging over time 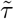, giving the exponential decay. We illustrate this in Figure 1(f) by showing *S*_**k**_(*τ*) for small Γ (small dash), intermediate Γ (solid line) and large Γ (large dash).

As stated earlier, the three aspects of adaptation, modulation and motion combine to produce the gained information. The product of the curves in Figures 1(e), (f) and (g) gives the three curves in Figure 1(h). Following (3), the integral of the curves in Figure 1(h) gives the information gain as encoded in *χ*_**k**_. Different aspects of the system dominate for different parameter choices. For example, when Γ is large the exponential decay of *S*_**k**_ in 1(f) is much faster than that the memory kernel autocorrelation in Figure 1(g) and so *S*_**k**_ is almost perfectly reproduced in the large Γ (large dash) curve in Figure 1(h). Conversely, when Γ is small, the slow decay of *S*_**k**_ in 1(f) means that the corresponding small Γ (small dash) curve in Figure 1(h) bears the imprint of the memory autocorrelation function. For all regimes the finite presentation time, as encoded by the autocorrelation function in 1(e), provides a cutoff for the curves in Figure 1(h). A particular case worth highlighting is that of very short presentation times (i.e. shorter than both the excitory response of the memory kernel, and the exponential decay of *S*_**k**_). Then, for a constant amplitude stimulus, we can approximate the gained information by

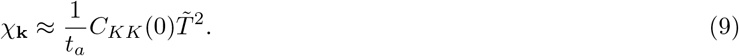

This quadratic dependence on presentation time will be relevant to our discussion of detection thresholds in Section IV.

In the remainder of this paper we first use the above model to consider contrast sensitivity at a specified wavevector for both constant and modulated stimuli. Then in Section IV we provide a more holistic view of detection thresholds, incorporating the sum over wavevectors present in most realistic stimulii. At certain points we present the consequences of our model with parameters appropriate for human vision. For the adaptation time and shape parameter we take *t*_*a*_ = 40 ms and 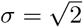, based on achieving approximate agreement between the autocorrelation of our memory kernel and that of a more involved one that has been shown to accord with the human visual system [38], see Section III of the Supplemental Material [45] for details. There is variability in the values given in the literature for the diffusion constant, and care is needed to ensure that microsaccades and head movements are not incorporated, but values of 0.01 − 0.04 arcmin^2^/ms have been reported [21, 47]. Based upon this, we take *D* = 0.02 arcmin^2^/ms unless otherwise stated. Lastly, we take blurring function *F* to have the form 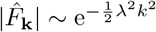 with *λ* ≈ 1 arcmin [48, 49].

**TABLE I.** A summary of the key notation used in this paper.

| Notation | Meaning |
| --- | --- |
| $t_a$ | *Adaptation time:* $\sqrt{t_s t_l}$, with $t_s$ and $t_l$ the short and long timescales of the memory kernel |
| $\sigma$ | *Shape parameter:* $\sqrt{t_l/t_s}$, with $t_s$ and $t_l$ the short and long timescales of the memory kernel |
| $\tilde{T}$ | *Dimensionless presentation time:* $T/t_a$, with $T$ the presentation time |
| $\Gamma$ | $k^2 D t_a$, with $2\pi/k$ a signal wavelength and $D$ the diffusion constant of the retinal motion |
| $\chi$ | *Measure of retinal information:* defined in (3) and surrounding text |

## III. EFFECTS OF RETINAL MOTION ON CONTRAST SENSITIVITY

With all the components in place, we now calculate explicitly the information gained from a stimulus presented at a particular wavevector, in terms of 𝒳_**k**_. The results of this section can also be viewed in terms of the contrast sensitivity as a function of the spatial frequency, |**k**|*/*2*π*, of a stimulus. The contrast sensitivity is proportional to 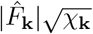, with 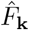 the Fourier transform of the blurring function. Therefore 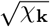 gives the contributions to contrast sensitivity from eye movement and signal modulation when passed through the temporal response. The contrast sensitivity function is a quintessential measure of the system-level performance of the visual system [50–54]. Although the classic experiments of Kelly [53] attempted to stabilise stimuli on the retina, it is now widely accepted that FEMs contribute to the contrast sensitivity function, and there have been a number of computational works investigating the role they play in this regard [1, 29, 55]. We consider first stimuli with constant amplitude, before moving onto sinusoidally modulated stimuli, for which the former is of course a limiting case.

### A. Constant amplitude

Taking our generic expression for 𝒳_**k**_ (3), substituting the particular forms of *C*_*KK*_, *C*_*AA*_ and *S*_**k**_ appropriate to a bi-exponential memory kernel, constant amplitude and diffusive retinal motion, and non-dimensionalising yields

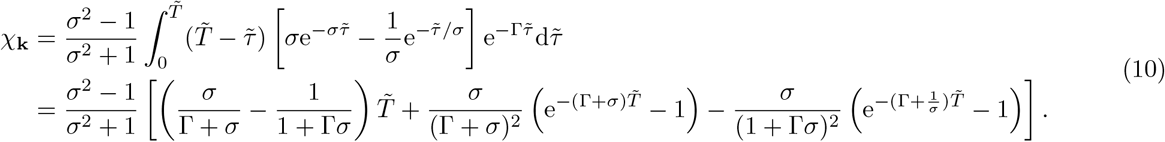

This is plotted as a function of Γ in Figure 2(a) for 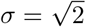 and a range of stimulus presentation times 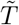, whilst in Figure 2(b) we show 𝒳_**k**_ as a function of 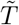 for several choices of Γ. There are several regimes of behaviour, which we now delineate. Figure 3(a) is a phase diagram indicating the values of Γ and 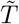 where each regime of behaviour applies. In what follows, it helps to define two constants that depend only upon the shape of the temporal response, parameterised via *σ*:

**FIG. 2.**
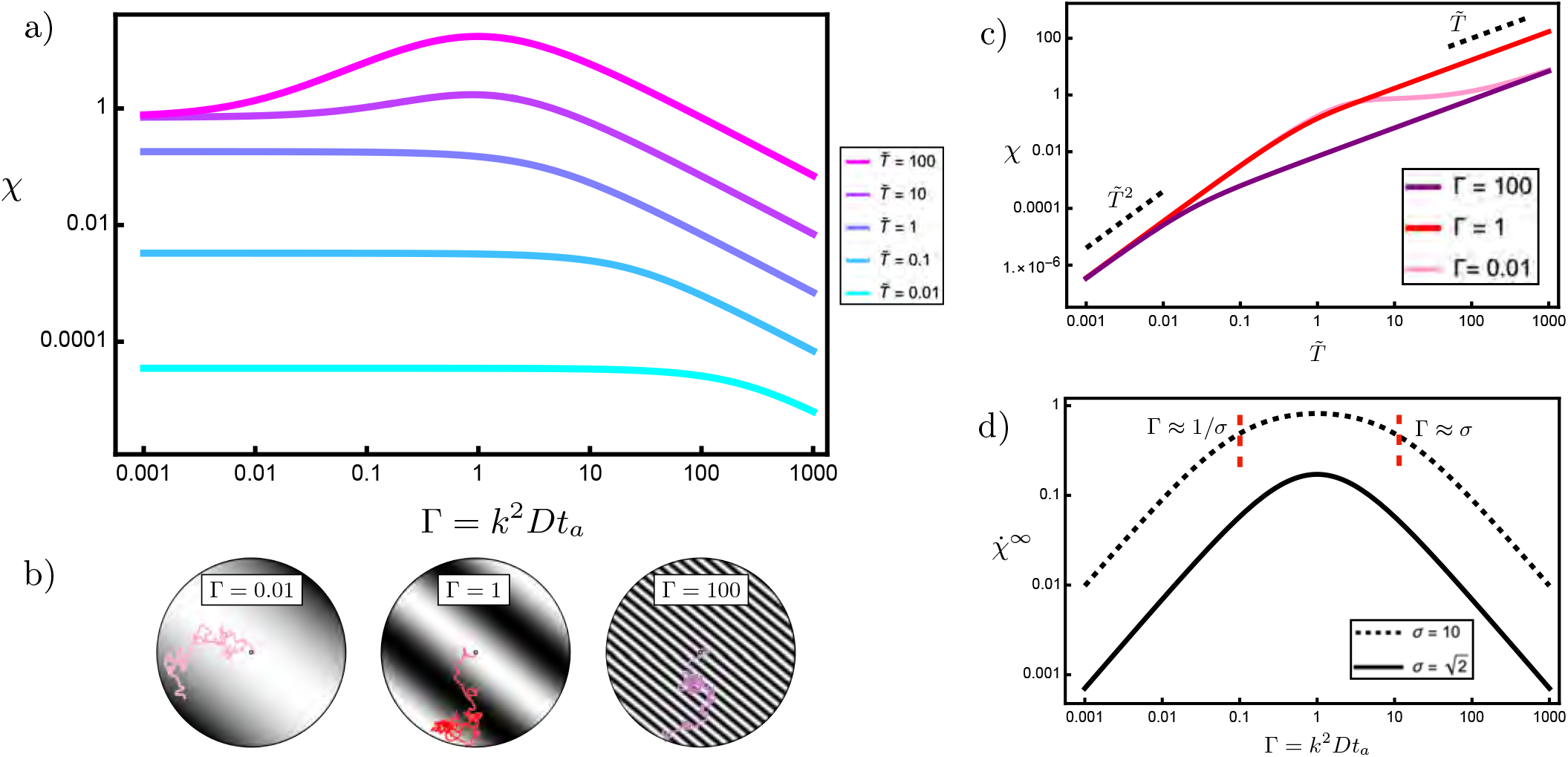
The retinal information gained from a constant amplitude signal with diffusive ocular drift. a) The information gained is captured by *χ*, given in general in (3) and in the particular case of a constant amplitude stimulus in (10). This is shown as a function of Γ, that is how far the eye diffuses in the adaptation time, measured in the wavelength of the signal (2*π/k*). The stimulus is presented for a time 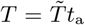, with 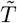 varying from 0.01 (cyan) to 100 (magenta) in powers of 10. The shape parameter, defined in terms of the long and short timescales of the memory function (*t*_*l*_ and *t*_*s*_) as 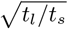, is 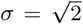.b) Schematic illustrations of different values of Γ, showing a grayscale stimulus with the path taken by the retina within the adaptation time. We emphasise that there is no absolute scale associated with these plots, and an increase in Γ = *k*^2^*Dt*_*a*_ may result from either increasing the spatial frequency in the stimulus, *k*, or increasing the diffusion constant, *D*. c) The information gained as a function of 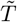 for three choices of Γ, highlighting the different power laws that arise. d) The asymptotic rate of information gain, 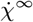. The solid curve corresponds to the same adaptive memory kernel as in a). The dashed curve illustrates the effect of a larger shape parameter 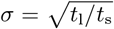, corresponding to a greater separation between the timescales of excitation and inhibition.

**FIG. 3.**
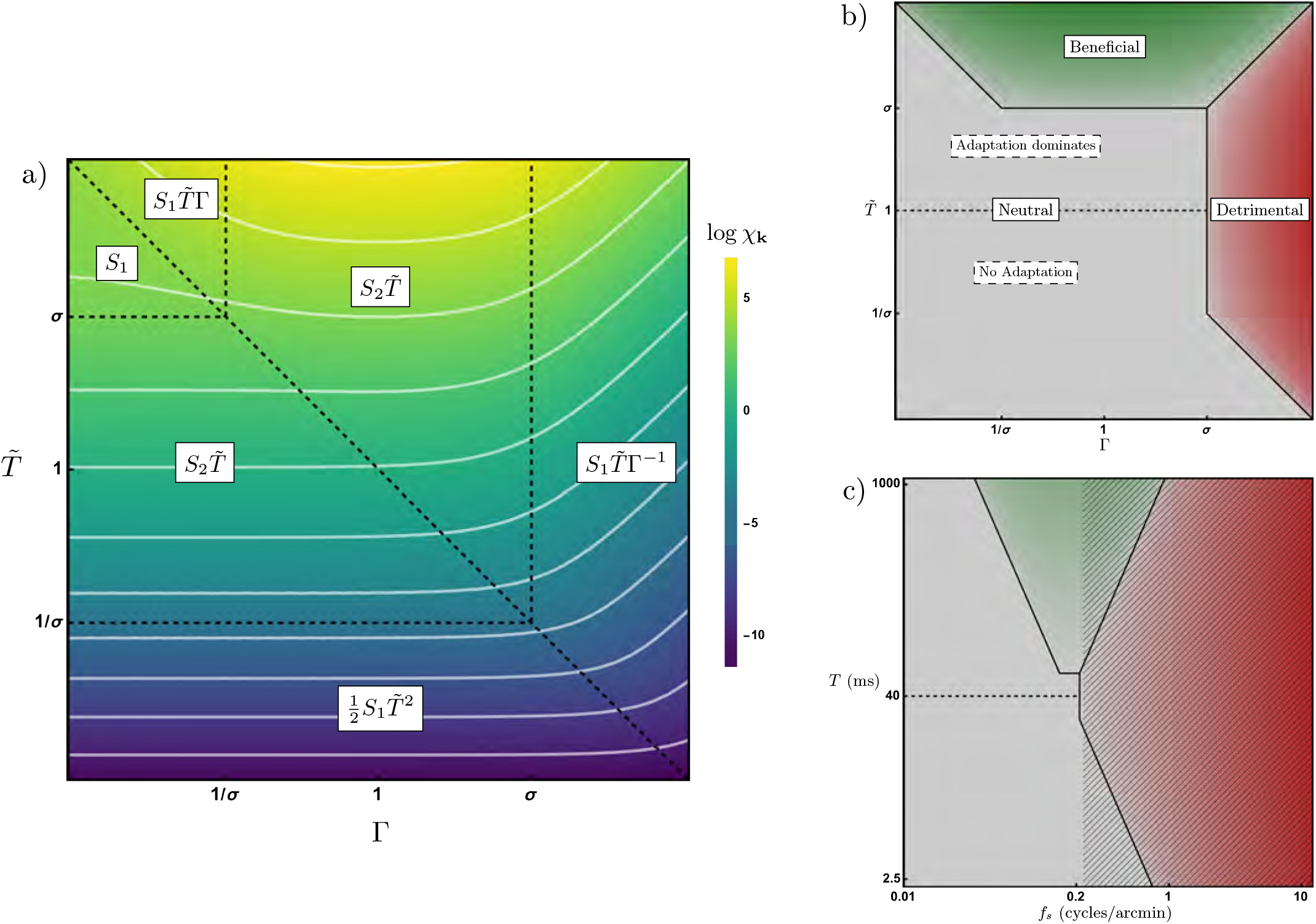
The phase diagram of fixational eye motion. a) The logarithm of 𝒳_**k**_, given in (10), as a function of Γ = *k*^2^*Dt*_*a*_ and the dimensionless presentation time 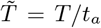. The white lines are contours of 𝒳_**k**_ and the labels indicate its behaviour in each region, with *S*_1_ and *S*_2_ the constants defined in (11) and (12). b) A simplified version of the same phase space as in panel a), indicating the regions in which eye motion is beneficial, detrimental or neutral in its effect. The dashed line at 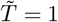 corresponds to the adaptation time. c) The eye motion phase diagram as appropriate for human vision, shown in terms of spatial frequency, *f*_*s*_, and presentation time *T*. In converting to dimensional units we have taken the adaptation time *t*_*a*_ = 40 ms (again indicated by a dashed line), the diffusion constant *D* = 0.02 arcmin^2^/ms and the shape parameter 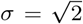. We have also assumed a blurring function of the form 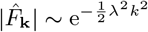with *λ ≈* 1 arcmin. The hatched region indicates the spatial frequencies for which information is lost due to this blur.

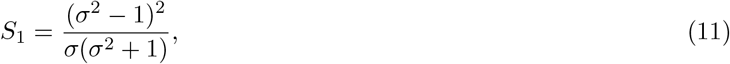

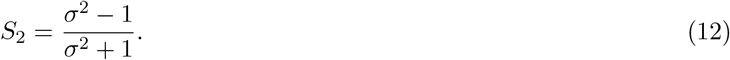

These two numbers appear in several of the subsequent expressions.

We note that for a stimulus at wavevector **k**, diffusive eye motion produces a temporal signal variation on timescale (*k*^2^*D*)^*−*1^. Comparing this with the presentation time *T*, we distinguish long presentation times (*T >* (*k*^2^*D*)^*−*1^, so that 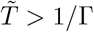) in which there is significant retinal motion over the spatial scale of the signal during stimulus presentation, and short presentation times 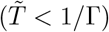 when retinal motion is small on the spatial scale of the signal during stimulus presentation.

For long presentation times, 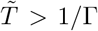, the first term of (10) dominates and the information gain grows linearly with time which is the behaviour seen in the upper right part of Figure 2(a) for large 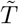. Schematic illustrations of different Γ values are shown in Figure 2b), and the corresponding time-dependence of 𝒳_**k**_ is shown in Figure 2c), with all curves exhibiting linear growth at large times. The resultant rate of information gain 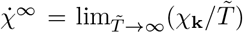 is shown in Figure 2(d), giving the shape of the peak in 𝒳_**k**_ that emerges at large presentation times. This depends on the extent to which the eye motion can drive information through the adaptive temporal response function. To assess this, we compare the signal variation timescale (*k*^2^*D*)^*−*1^ with the two timescales in the memory kernel for retinal response, *t*_l_ and *t*_s_. This gives rise to three regimes of behaviour: (i) the temporal signal variation is slower than the inhibitory component of retinal response (*k*^2^*Dt*_l_ ≪ 1 so that Γ ≪ 1*/σ*), (ii) temporal variation is slower than the excitory response but faster than the inhibitory component (*k*^2^*Dt*_l_ ≫ 1 but *k*^2^*Dt*_s_ ≪ 1 so that 1*/σ* ≪ Γ ≪ *σ*), or (iii) temporal variation is faster than both excitory and inhibitory response (*k*^2^*Dt*_s_ ≪ 1 so that Γ ≫*σ*). We briefly describe the rate of information gain in each of these regimes.

When Γ ≪ 1*/σ* the signal variation at a receptor due to eye motion is slower than any component of the temporal response function. In the absence of any eye movement, when Γ = 0, adaptation is perfect and there is no component of 𝒳_**k**_ proportional to 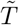, meaning increasing the presentation time has no further effect on the transmitted information. For small non-zero Γ, the eye movement marginally offsets the adaptation, so that some signal is transmitted through the temporal filter. The extent to which this occurs is proportional to Γ, so that

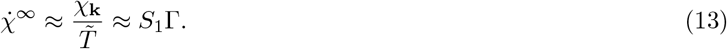

This appears in Figure 2(d) as a gradient of one.

For 1*/σ* ≪ Γ ≪ *σ*, the short-time initial exponential decay of the temporal response dominates, yet the eye motion decorrelates the signal before the negative, adaptive part of the temporal response can take effect, hence the eye movement overcomes adaptation. Within this regime the rate becomes independent of Γ, with

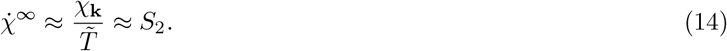

For this regime to be well-defined, we must have *σ* ≫ 1, that is the short and long timescales of the temporal filter, *t*_*s*_ and *t*_*l*_, must be well separated. We illustrate this in Figure 2(d) by showing the rate of information gain for *σ* = 10, so that we can see an intermediate plateau beginning to open up between Γ = 1*/σ* and Γ = *σ*, for which 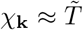. Physiologically, it seems unlikely that there is such strong separation in the temporal filter [38, 46], and so this intermediate plateau is not so well defined, as is clear when 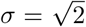. In practice (14) then represents an upper bound of the degree to which eye motion can overcome adaptation.

Lastly, when Γ ≫ *σ* the retinal motion is so rapid as to vary the incoming signal faster than the short-time positive component of the temporal response. Hence the temporal response becomes irrelevant and the result depends only on the signal decorrelation due to eye motion, so that 𝒳_**k**_ decreases with Γ as

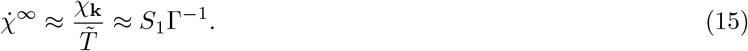

This appears as a gradient of minus one in Figure 2(d).

In summary, for long presentation times 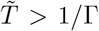 the rate of information gain is proportional to Γ for small Γ, proportional to 1*/*Γ for large Γ and, if the short and long time components of the temporal response are well separated, constant for intermediate values of Γ. This large 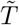 limit for the contrast sensitivity, exhibiting a peak at a given spatial wavenumber, has been described in computational work [29, 47, 55], and is consistent with experimental observation of contrast sensitivity functions [46].

We now consider how this picture is changed by shorter presentation times, 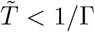, paying particular attention to the dependence of 𝒳_**k**_ on 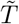, since, as highlighted at the end of Section II B, this relationship will not always be linear. For short presentation times there is no significant retinal motion on the spatial scale of the signal and the response due to switching the stimulus on and off dominates over those induced by eye motion. Hence, the retinal motion can be neglected and the information gained becomes independent of Γ, leading to the horizontal lines on the left of Figure 2a), the heights of which depend only on the presentation time. Focusing on this short presentation time limit, there are three sub-regimes, which, in analogy with those for Γ presented above, are defined by whether the presentation time is shorter than both adaptive timescales 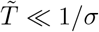, intermediate to them, 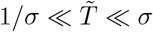, or longer than both, 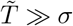. We find that these three regimes cause the information gained to first increase quadratically with presentation time, then increase linearly and finally stagnate to a constant value, as we now briefly explain.

When the presentation time is shorter than the short-time response of the adaptive filter, the stimulus is being integrated such that the transmitted signal grows linearly with presentation time. Since 𝒳_**k**_ depends on the square of the transmitted signal, it grows quadratically with presentation time

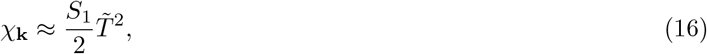

displayed by all curves in 2c), this being the scenario highlighted at the end of Section II B. If the adaptive shape parameter is sufficiently large then there will a regime where the presentation time is between the short and long adaptive timescales, meaning the signal disappears before the negative, adaptive part of the temporal response can take effect. This is analogous to the situation yielding (14), and so again 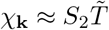. It is not seen in 2c), since for all these curves 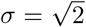. Finally, when 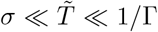 (which requires that Γ ≪ 1*/σ*) the gained information saturates to a constant value given by

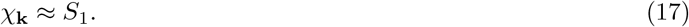

This saturation of information gain is due to the observation time being long enough for adaptation to occur, but not long enough for the retinal motion to circumvent this adaptation by changing the signal received by the photoreceptors. This can be observed for the pink curve in 2c), this being the only one with a small enough value of Γ to allow this regime to arise. As expected, the information plateau spans 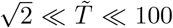, with the asymptotic information rate discussed earlier emerging for 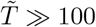, in this case given by (13).

As noted above, Figure 3a) presents a phase plot indicating the values of Γ and 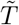 where each regime of behaviour in 𝒳_**k**_ applies. This delineates the region for 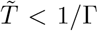, where the response is dominated by transients arising from temporal modulation of the external stimulus via *A*(*t*), and the region for 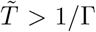 where the eye motion begins to have significant effect on information gained, and where a peak in 𝒳_**k**_ emerges at Γ = 1.

Having determined the behaviour of 𝒳_**k**_ across this phase space, we can naturally establish the regions in which fixational eye movements have a beneficial, detrimental or neutral effect on the acquisition of visual information, as shown in Figure 3b). To construct this diagram, at each presentation time 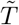, we compare *χ*_**k**_ with its value 𝒳_**k**,Γ=0_ at Γ = 0 which corresponds to zero eye motion. When 𝒳_**k**_ *>* 𝒳_**k**,Γ=0_ then eye motion can be said to enhance the transmitted information, and so is beneficial. When 𝒳_**k**_ *<* 𝒳_**k**,Γ=0_ then eye motion is detrimental to transmitted signal, whilst equality 𝒳_**k**_ = 𝒳_**k**,Γ=0_ implies a neutral effect. This procedure gives rise to the regions sketched in Figure 3b). Given the long-running debate as to whether fixational eye movements have a negative, blurring, impact [8, 56] or usefully modify the spatio-temporal structure of the stimulus [18, 29–32, 34, 57, 58], it is striking that all three of these regions exist.

To understand what this phase diagram implies for human vision, we convert to dimensional quantities, taking the adaptation time *t*_*a*_ = 40 ms, the diffusion constant *D* = 0.02 arcmin^2^/ms and the shape parameter 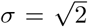. This results in the phase diagram shown in Figure 3c) in terms of stimulus spatial frequency, *f*_*s*_, in cycles/arcmin and presentation time in ms. We have also assumed a blurring function of the form 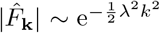, with λ ≈ 1 arcmin. Taking 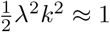 as the criterion for blur having a significant effect, we find that blur eliminates the information at high spatial frequencies with *f*_*s*_ ≳ 0.2 cycles/arcmin, this being the hatched region in Figure 3c).

The practical consequence of the above is that for presentation times that are long compared to the adaptive timescales, that is *T* ≳ 50 ms with the assumed parameter values, corresponding to 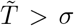, there is clear benefit from eye motion within the green portion of Figures 3b) and c), with a pronounced peak in the gained information emerging at Γ = 1 for the magenta curves in Figure 2a) and an associated hotspot in Figure 3a). The height of the peak is proportional to the presentation time, with a shape given by (13)-(15). Once again, when 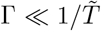 the eye motion becomes insignificant and the gain of information saturates due to adaptation, according to (17). The overall shape of the emergent peak is consistent with the prior numerical analysis [29, 47, 55]. Not only do we reproduce this peak, but our theory provides a simple prediction for where it occurs. The peak exists for long presentation times because, once adaptation has set in, it is possible to have retinal motion which is neither irrelevant, nor purely decohering, but rather of appropriate magnitude to offset the adaptive effects. This is at the heart of the peak’s location. It occurs when Γ = 1, that is when the retinal motion induces temporal variation matched to the frequency of the band-pass adaptive filter. It should be highlighted that, as per our definition of Γ, the optimal eye motion is inherently stimulus-dependent, since the induced time-variation is a product of both the eye’s motion and the spatial variation of the stimulus. This will be important for our discussion of detection thresholds in IV.

On the other hand, for presentation times shorter than ∼ 50 ms, that is 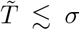 with our choice of physical parameters, eye motion can only be deleterious towards signal perception. This is illustrated by the cyan curve in Figure 2a. On such timescales adaptation is incomplete, and so the retinal motion is either insignificant or simply impeding the integration of the signal. The former produces the horizontal lines and flat contours to the left of Figure 2a) and Figure 3a) respectively, with the gained information following (14) or (16). In the latter case, information gain falls according to (15), leading to the gradients of minus one on the right of Figure 2a), and correspondingly gradients of one in the contours to the right of Figure 3a). This decrease occurs once the retinal motion is significant over the presentation window, 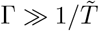 and dominates the adaptive timescales Γ ≫ *σ*. Again, this pair of requirements means that for short presentations the eye motion is either irrelevant or dominating the decoherence of the signal; there is no regime in which they provide a benefit. This is made apparent in the red portion of Figures 3b) and c), and can also be seen in the monotonic behaviour of the contours in Figure 3a) for 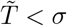 and the lower three curves in Figure 2a), for which the same condition holds.

However, this fall off in signal perception may not be of great consequence if it occurs for spatial wavelengths which are already suppressed due to optical blur or finite size of receptors, incorporated in our description through the blurring function *F* in (1). As noted above, we take this blurring function to have the form 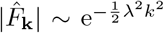, with *λ* ≈ 1 arcmin. As shown in Figure 3c), this means that with parameter values appropriate to human vision, the region in which eye motion is detrimental is essentially covered by the region in which information is already lost due to blur. This means that the magnitude of diffusion observed in human vision [21, 47] is optimal to provide infomational benefit, given the presence of optical blur and hints at another instance of biological systems achieving physically imposed performance limits [9, 10].

### B. Sinusoidal modulation

We now consider a stimulus whose amplitude is modulated sinusoidally, such that *A*(*t*) = cos(*ωt*). The resulting behaviour depends on how the oscillation frequency compares to the adaptation time, so we find it convenient to define a non-dimensional oscillation frequency 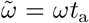, and to consider a range of values of 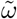. Figure 4 shows the impact of modulation frequency on the information gain. The corresponding expression for 𝒳_**k**_ and its derivation is given in Section V of the Supplemental Material [45]. For very short presentation times, so that 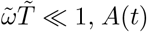 is approximately constant during the presentation time and so the results match those of the previous section. Otherwise, there are several key differences compared to the constant amplitude case. One minor difference is that the information gained does not always increase monotonically with presentation time 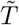, since an alternating positive and negative signal can cancel when passed through the temporal filter [38]. Typically the non-monotonic behaviour occurs for small Γ, where retinal motion is insignificant, and at 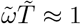, where the effects of alternating signal become apparent, as seen e.g. in Figure 4(a) and (c).

**FIG. 4.**
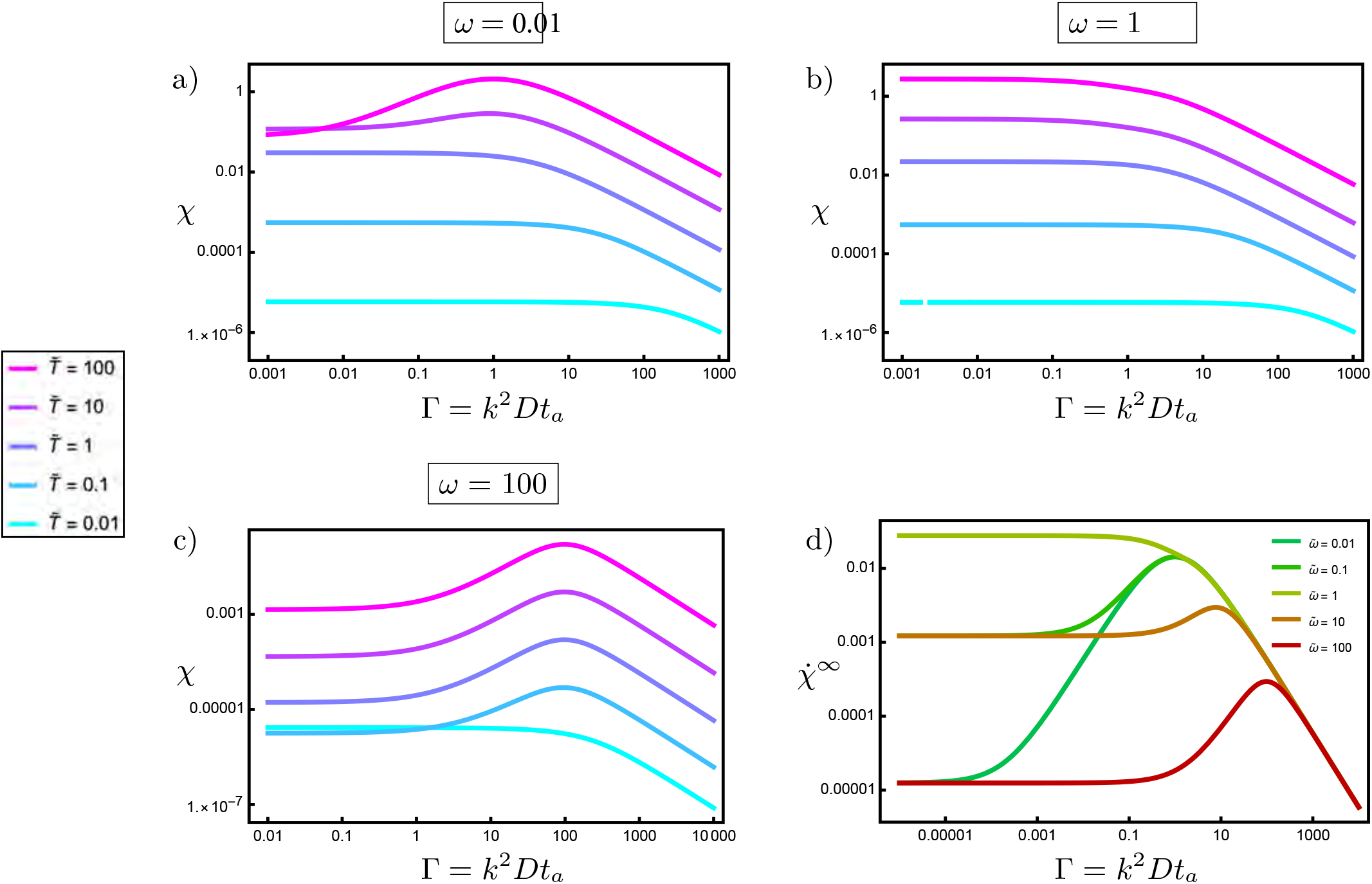
The retinal information gained from a signal modulated sinusoidally at frequency 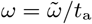 with diffusive ocular drift. a)-c) For the coloured curves 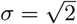 and 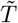 varies from 0.01 (cyan) to 100 (magenta) in powers of 10. d) The asymptotic rates of information gain for 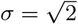 and 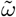 varying from 0.01 (green) to 100 (red) in powers of 10.

It is helpful to first consider the effect of sinusoidal modulation in the absence of eye motion (Γ = 0), before discussing how eye motion perturbs the result. Here, the signal modulation provides a non-constant signal to the temporal response, so that perfect adaptation no longer occurs. In the limit of long presentation times, 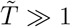, this gives rise to a constant rate of information acquisition, so that:

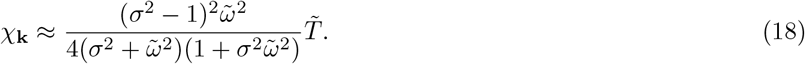

This is maximised at 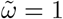 (and is almost constant for 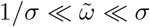), indicating that the transmission of information through the temporal response is optimised when the oscillation frequency is matched to the adaptation time. In this case, the oscillations of *C*_*AA*_ in Figure 3 from the Supplemental Material [45] match well the positive and negative portions of *C*_*KK*_ in Figure 1(g). Then, given that the signal modulation is optimally overcoming adaptation, adding eye motion can only make things worse. Hence, in figure 4(b), 𝒳_**k**_ is always a decreasing function of Γ, transitioning from a constant value towards 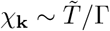 above a critical value of Γ (which is Γ = 1 for 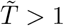, or 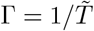 for 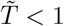).

For smaller frequencies 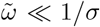, the modulation is too slow to optimally overcome adaptation. In this case eye movement still helps, so that a peak in 𝒳_**k**_ at Γ = 1 emerges when 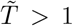, as can be seen in Figure 4(a). This is qualitatively similar to the constant amplitude case in the previous section, except that in the limit of low eye movement, Γ → 0, the plateau in *χ*_**k**_ continues to increase when 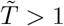 because of the signal modulation.

Finally, for high oscillation frequencies, 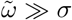, the signal modulation is more rapid than the integration time of the fast component of the temporal response. Hence, the temporal response averages over the modulated signal, giving a low output in the absence of eye movement. In this case, eye movement can help, not by overcoming adaptation, but rather by spatially decorrelating the signal faster than the amplitude modulation. This occurs when the decay of *S*_**k**_ in Figure 1(f) matches well the oscillations of *C*_*AA*_ in Figure 3 from the Supplemental Material [45], cutting out the negative portion of *C*_*AA*_. The optimum is when the rate of eye movement matches the signal modulation, giving a peak in 𝒳_**k**_ at 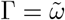 which emerges for 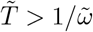, as illustrated in Figure 4c).

Hence, for long presentation times 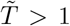 we find that eye motion helps both when 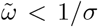 (when the motion continues to help overcome adaptation) and for 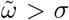 (when the eye movement offsets the signal modulation). For 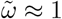 the eye movement is no longer beneficial. With eye motion now included, the long time behaviour is given by:

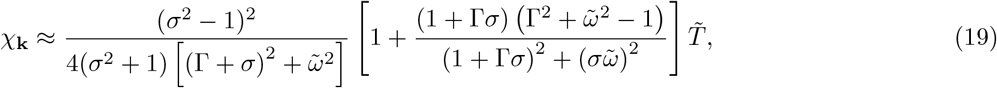

the corresponding rates being the curves shown in Figure 4(d). Recalling that the contrast sensitivity is proportional to 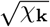, Eq. (19) and Figure 4(d) can be considered to represent the contributions of eye movement, signal modulation and temporal response to the spatiotemporal contrast sensitivity function [50–53, 55]. The symmetry between a frequency and its inverse for small Γ is a consequence of our choice of temporal response function, although the power laws for small and large frequencies are generic. A similar symmetry is apparent in Figure 2(b).

The full spatiotemporal contrast sensitivity function, CSF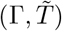 is attained by incorporating blur, which, as in the construction of Figure 3c), we take to be form the form 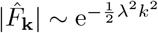, giving

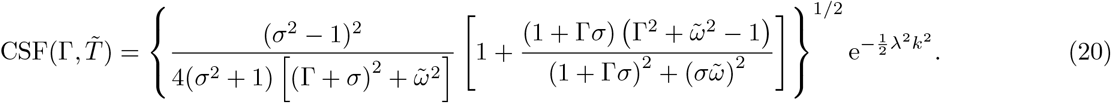

This is shown in Figure 5 as a function of spatial frequency, *f*_*s*_, in cycles/arcmin and temporal frequency, *f*_*t*_, in cycles/ms. In converting to dimensional units we have used the same parameter values as in Figure 3c), namely *t*_*a*_ = 40 ms, 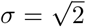 and *λ* = 1 arcmin. The case with no eye motion, *D* = 0, is shown in Figure 5a) and exhibits a band-pass response in temporal frequency, owing to the adaptive linear response, and low-pass behaviour in spatial frequency, due to blur.

**FIG. 5.**
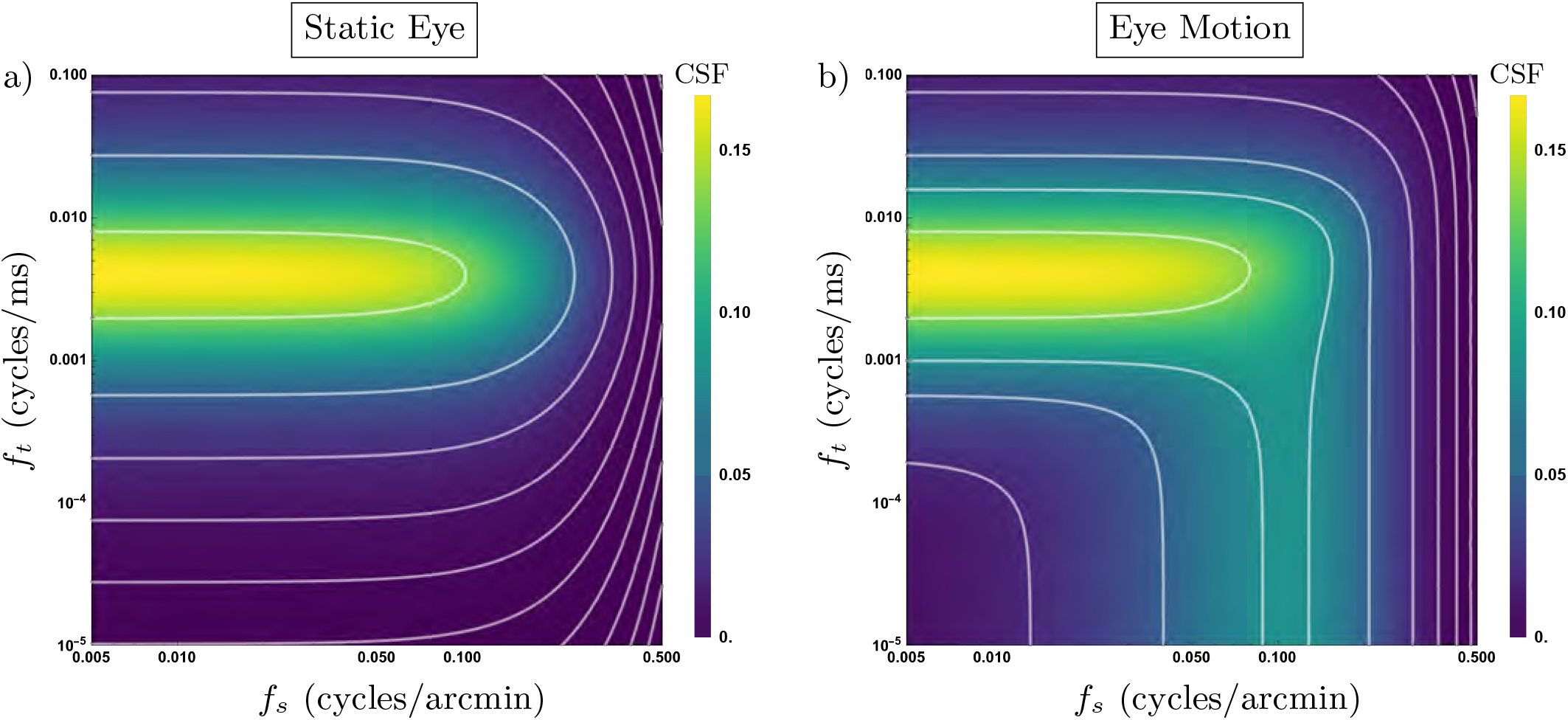
Spatiotemporal contrast sensitivity functions. The contrast sensitivity function, given in (20) is presented as a function of spatial and temporal stimulus frequency, *f*_*s*_ and *f*_*t*_ for no eye motion, a),and *D* = 0.02 arcmin^2^/ms, b). In both cases theadaptation time has been taken as *t*_*a*_ = 40 ms, the shape parameter as 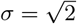 and the blur lengthscale as *λ* = 1 arcmin. The white lines correspond to contours of the contrast sensitivity function.

The case with eye motion is illustrated in Figure 5b) for *D* = 0.02 arcmin^2^/ms. The contrast sensitivity function is now band-pass in both temporal and spatial frequencies. When the temporal frequency is below its band-pass value, the optimal spatial frequency is constant. However, for higher temporal frequencies, the optimal spatial frequency is shifted upwards, as indicated by the outwards bowing contour in Figure 5b). This behaviour mirrors that shown in Figure 4d), where 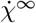 peaks at Γ = 1 for 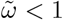, but for 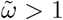 the peak moves to the right, occurring at 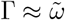.

We have considered the contrast sensitivity function in the limit of infinite presentation time, that is, using 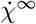, but we can infer the effect of finite presentation times from Figure 4a)-c). Reducing the presentation time flattens the rise in *χ*_**k**_, such that the peak is diminished relative to its value for low Γ. In the context of Figure 5 this corresponds to filling in the region where both *f*_*s*_ and *f*_*t*_ are low. For sufficiently short presentation times the band-pass spatial response vanishes, as indicated by the cyan curves in Figure 4a)-c).

The key message from Figure 5 is that, by converting spatial variation in a stimulus into temporal variation in the photoreceptor signal, eye movements can transmute the adaptive band-pass response for temporal frequencies into a band-pass response for spatial frequencies. Even though it has been established that retinal ganglion cells also produce a spatial bandpass effect, [59–63], it is striking that such mechanisms are not required to reproduce the features of the spatiotemporal contrast sensitivity function, this can be done with eye motion alone.

With this eye-motion-mediated connection between temporal and spatial adaptation in hand, we are now in a position to consider whether the apparent scale-dependent adaptation seen in experiments of Barlow [37] may also be explained by eye movements.

## IV. DETECTION THRESHOLDS

In the previous section we determined the impact of retinal motion on the eye’s ability to gain information from a signal with a given spatial frequency. We now extend this to consider the impact of this movement on the threshold amplitude for detection of a target, in which many spatial frequencies are superposed. In doing this we employ a model of detection probability, *p*, proposed by Watson [38]. The underlying principle of the model is that detection of a stimulus requires only that the stimulus be detected above the background by at least one of the available receptors, at one or more times during the observation window. As shown in Section I D of the Supplemental Material [45], this model gives a detection probability of

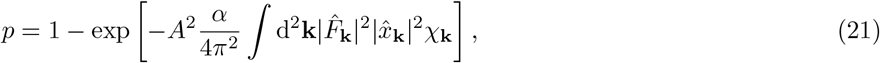

where *α* is a normalisation constant and we have described the stimulus in terms of a spatial form 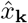 and an overall intensity *A*. The stimulus intensity *A* can be varied to find the threshold for detection, which, from (21), is

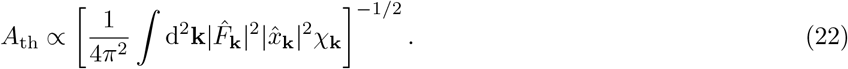

We consider the experimental paradigm employed by Barlow [37], in which detection thresholds were measured for circular stimuli of two different sizes: a large target of radius 177 arcmin and a small one of radius 3.5 arcmin. For both targets it was found that the threshold intensity initially decreased with the inverse of the presentation time. For large stimuli the threshold intensity then did not change with further increases in presentation time, due to adaptation. However, for a small stimulus the threshold intensity continued to decrease, but at a reduced rate.

To explain this behaviour, it has been suggested that the intrinsic retinal response has a form of temporal adaptation with some spatial dependence, such that low spatial frequencies are perfectly adapted, but high spatial frequencies are imperfectly adapted [38]. Such a retinal response cannot be modelled within the simple framework presented in this paper, as presented in Section II A, using a separate temporal response function *K* and spatial blurring function *F*. Instead, this explanation requires that the functions *K* and *F* should be combined into a non-factorisable spatiotemporal retinal response function (i.e. this explanation requires that the intrinsic spatiotemporal contrast sensitivity function of the retina cannot be separated into the product of spatial and temporal functions). It is certainly possible that the retinal ganglion cells produce such a complex spatiotemporal response, for example by giving rise to multiple channels of spatial response, each with their own temporal behaviour [38, 64]. However, we shall show that all features of the experiments can be explained without invoking such scale-dependent adaptation. The necessary spatiotemporal complexity arises naturally within our minimal model, via the interaction of retinal motion with an adaptive temporal response function.

For a signal with constant intensity within a circle of radius *R*, and zero outside this, a standard calculation gives 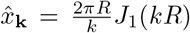, with *J*_1_ a Bessel function of the first kind. Additionally we once again assume an exponential blurring function, 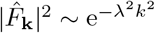. Consequently,

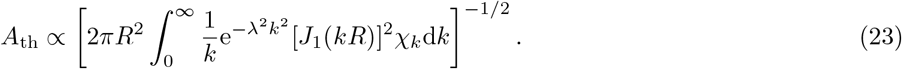

The integral is a sum over 𝒳_**k**_, the rate of gaining information from a wavevector **k**, weighted by the prevalence of that wavevector in the signal and the blurring function. In Figure 6 we plot this threshold intensity *A*_th_ as a function of stimulus presentation time *T*, both for (a) the large target of radius 177 arcmin and (b) the small one of radius 3.5 arcmin. Here we use the same parameterisation as in previous figures, i.e. adaptation time *t*_*a*_ = 40 ms, shape parameter 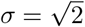 and blur lengthscale *λ* = 1 arcmin, but we show results for a range of diffusion constants *D* from 0.00125 min^2^*/*ms (cyan) to 0.04 min^2^*/*ms (magenta).

**FIG. 6.**
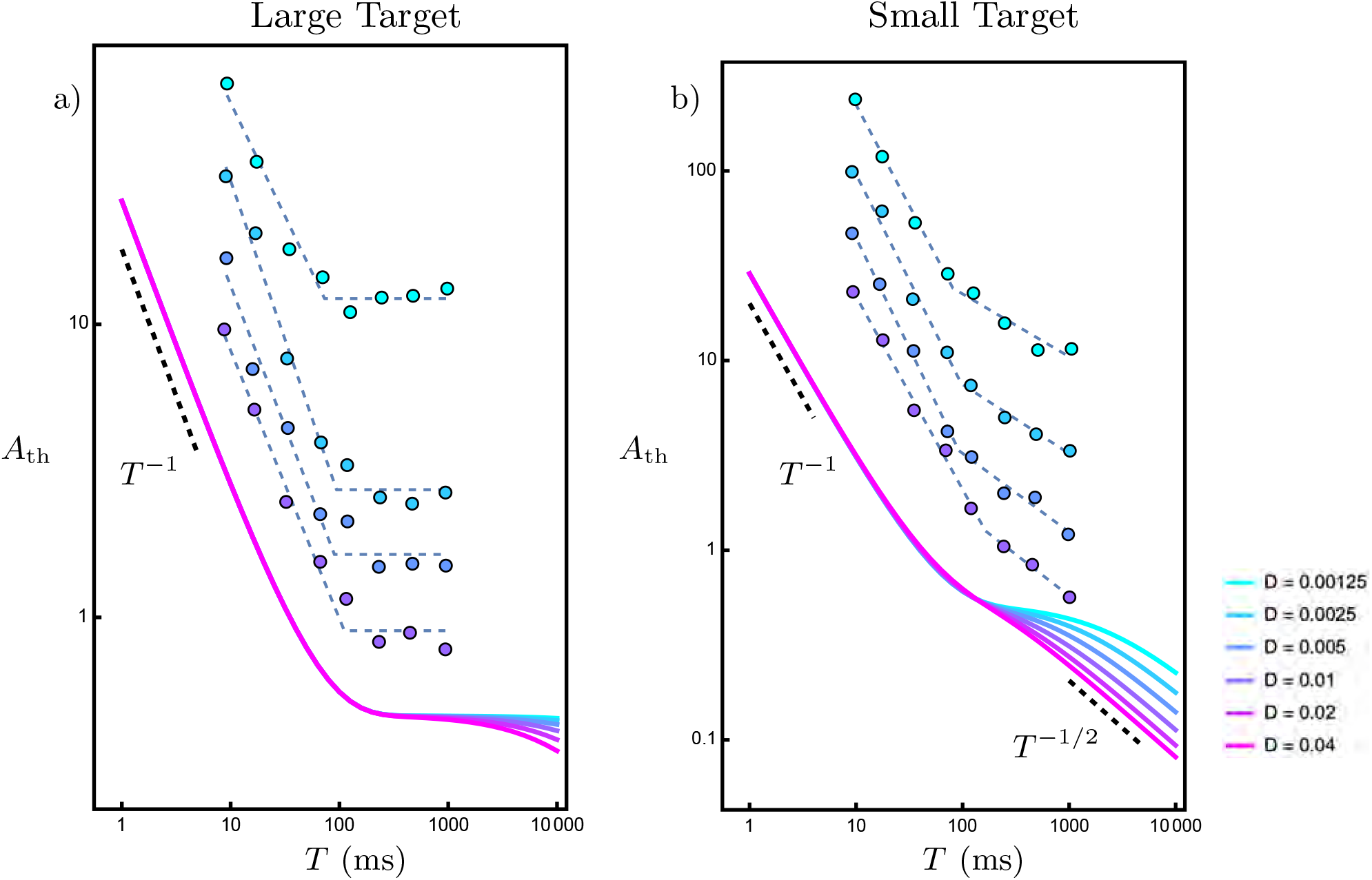
The contrast detection threshold, *A*_th_, for a circular stimulus of radius *R* (measured in arcmin) as a function of observation time, *T*, (in ms). The dots show the data from Barlow’s experiments [37], with the dashed lines added to guide the eye, and the solid lines the prediction of (23) for a) the large target (*R* = 177 arcmin) and b) the small target (*R* = 3.5 arcmin). The occular drift is taken to be diffusive with diffusion constant *D* (measured in min^2^*/*ms). In both panels *D* is varied in powers of 2 from 0.00125 min^2^*/*ms (cyan) to 0.04 min^2^*/*ms (magenta). The data from Barlow’s experiments have been coloured according to the background intensity, the logarithm of which is 7.83 (cyan), 5.94, 4.96 and 3.65 (purple). Since our theory predicts only the dependence of *A*_th_ with *T*, not its absolute value, we have applied an overall shift to the datasets to ease presentation.

Although there does not exist a closed form for *A*_th_, we can understand all its essential features from a few simple arguments. As shown in III A, at short times 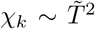, while at long times 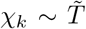, in both cases for all *k*. This immediately produces the short time 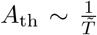 behaviour seen in [37], as well as a 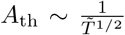 dependence at long times. Both these regimes are indicated with black dashed lines in Figure 6. At intermediate times there is a plateau due to adaptation causing the gain of information to stagnate. In III A we showed that this stagnation begins after full adaptation, when 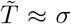, i.e. *T* = *t*_*l*_, the long adaptation time (of order 60 ms for our parameterisation). This stagnation is apparent in both Figure 6(a) and (b), as the time at which the plateau commences. The information stagnation ends once there has been significant retinal motion to overcome adaptation, that is when 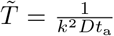, i.e. 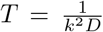. Hence, irrespective of the stimulus size, we expect the plateau to shorten as the diffusion constant *D* increases, which is again apparent in both Figure 6(a) and (b).

The crucial point is that a given eye motion does not affect all spatial scales in the stimulus equally, and this inherent scale-dependence replaces the need for any scale-dependence in the form of adaptation. Making the target smaller, i.e. decreasing *R*, puts the relevant visual information at smaller length scales, or larger spatial frequencies *k*. Hence 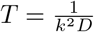 is smaller at the relevant scale and so decreasing *R* shortens the plateau, as is clear when comparing Figures 6(a) and (b). Indeed for the small stimulus size of radius 3.5 arcmin in Figure 6(b), when *D* is sufficiently large (0.01 arcmin^2^/ms or larger) the plateau almost entirely vanishes, so that one finds only a transition region of indeterminate slope between the regimes of 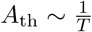 and 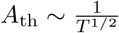.

Figures 6(a) and (b) also show the results of Barlow [37]. In these experiments the contrast threshold for the large and small stimulii were measured for various background light intensities, shown as separate datasets on the plots. We have ommitted data from the lowest light intensity, since visual response is known to be altered at low intensities [46, 65, 66]. Since our theory predicts only the dependence of *A*_th_ with *T*, not its absolute value, we have applied an overall shift to the datasets to ease presentation. It is clear that with a reasonable choice of diffusion constant (say *D* = 0.02 arcmin^2^/s), and with the reasonable choices for other parameters described above, our minimal model matches the temporal behaviour of the experiments extremely well. In particular, we reproduce the central result of stagnation of contrast detection threshold for the large stimulus, but that the contrast detection threshold continues to decrease for the smaller stimulus.

In practice, it would be impractical to run sufficiently long experiments to see the fully established 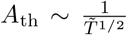 regime, and indeed additional physiology may obscure it. Hence the time axis in Figure 6(a) is truncated in the experiments by Barlow, and we believe that what was observed is perfect adaptation for large targets, and the transition towards the 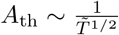 regime for smaller targets.

The experimental observations of the qualitative difference in contrast threshold with stimulus size can thus be explained solely via the space-time coupling produced by fixational eye movements. These effects are seen even within our minimal model. It is thus not necessary to invoke complex lengthscale dependence of the temporal adaptive response to explain these data. However, as noted above, such conjectures exist in the literature [38, 64]. This points to the need for additional experiments that would tease apart these two possibilities. A suitable investigation could involve comparing constrast detection thresholds as a function of presentation time, for stimuli that are either stabilized or unstabilized on the retina. Here, if the effects observed by Barlow are accounted for by eye movements, then with the effects of eye movement removed via stabilisation the temporal response to small stimuli should more closely match those of large stimuli.

## V. DISCUSSION

We have formulated an analytic framework for determining the impact of ocular drift on the information acquired from a stimulus. Our measure of the acquired information may be derived from the mutual information between the stimulus and a minimal model for the response of the early visual system, which incorporates drift, along with blur and a band-pass temporal response. Taking the drift to be diffusive, we identify the key dimensionless parameters which underpin the interactions between the stimulus structure, temporal response and ocular drift. These interactions are phenomenologically rich, as we summarise in a phase diagram, and have the consequence that ocular drift may be beneficial, detrimental or negligible in its impact on the acquisition of visual information. We find that optimal performance is achieved when *k*^2^*Dt*_*a*_ ≈ 1, where *k* is the wavevector of the stimulus, *D* is the diffusion constant of the drift and *t*_*a*_ is a timescale characterising the adaptive temporal response.

Working with parameters appropriate for human vision, our model reveals that the stimulus wavevector for which the observed degree of drift has the greatest benefit and the wavevector above which information is lost due to blur are the same. This suggests that, far from merely being noise, ocular drift is tuned to maximise the gain of visual information given the constraint of optical blur. Although such a statement exists in the literature [29], our work shows concretely how this arises from different components of the model. Additionally, a key emergent principle is that retinal motion combined with a band-pass temporal response produces an effective band-pass spatial response, even though such a response is not an explicit component of the model. This is shown most fundamentally in the modelled contrast sensitivity function. Although the existence of neural spatial receptive fields with a so-called centre-surround (bi-phasic) structure is well established [59–63], we believe it important to note that the data at the system level can be qualitatively explained without this. Furthermore, the model provides an alternative qualitative explanation for Barlow’s data on threshold detection amplitudes for stimuli of different sizes as a function of time [37]. These data have been taken as evidence for multiple spatial mechanisms that differ in their temporal response, such that smaller stimuli are postulated to recruit detection mechanisms with more sustained temporal impulse response, meaning that the integral of the linear temporal response function is not zero, but positive [38]. The model instead shows that the observed behaviour can emerge from the combination of eye-movements and a biphasic temporal response. Indeed, our work indicates a need for further experiments, repeating those of Barlow but with the stimulus stabilised on the retina, to distinguish the roles that ocular drift and spatio-temporal tuning of neural mechansisms play in producing the observed behaviour.

There are various ways in which our calculational approach could be extended. For example, one could relax the condition that the spatial and temporal variation of stimulus be separable, or consider a prior distribution of stimuli consistent with the statistics of natural images [67–69]. In this latter context it would be interesting to turn the calculation on its head, such that rather than calculating the effect of a prescribed ensemble of retinal motion on the available visual information, we instead determine the optimal form of ocular drift on the basis that optimality corresponds to whitening the power spectrum [1, 29, 41, 70].

In this paper we have taken a minimal approach to physiological details, both by neglecting certain features and in the modelling of those we retain. However, our framework is sufficiently generic to allow more detail to be incorporated as desired. There is no need to take the form of ocular drift to be diffusive and indeed, although not explored here, our theory indicates that a degree of persistence may be beneficial. While we have taken the spatial response to be a low-pass filter originating from optical blur, there is no obstruction to incorporating a band-pass spatial response. It would then be interesting to study both separable and non-separable forms of spatial-temporal response, particular given the latter’s relevance to attempts to connect Barlow’s data [37] to postulated underlying physiological mechanisms, as described above. We have taken the photoreceptor array to be continuous and homogeneous, neglecting, amongst other things, the discrete nature of photoreceptors and the high-acuity foveola in the centre of the retina. Incorporating these features into our model would provide insight into other hypotheses for the function of FEMs, such that they act to maintain a prefered retinal location [71], or achieve super-resolution through appropriate sampling of an image [72, 73].

Nonetheless, we maintain that this work does not suffer by omitting such details. This allows us to expose the basic principles underpinning the function of ocular drift in vision. Moreover, it lets us trade specificity for universality and see that, while the effects we have described apply to vision, they are not limited to that context. Motion and some form of adaptive response to the environment are near-ubiquitous signatures of life and our results indicate how these may interact non-trivially when an organism tries to learn about its surroundings. As one example, bacteria employ a band-pass filter to detect chemical gradients, which they traverse via runs interrupted by stochastic tumbles. Though the mechanisms behind the two are different, one might wonder whether these tumblings have a similar impact on sensing chemical signals as ocular drift has on detecting visual ones.

## Acknowledgements

This work was supported by the Engineering and Physical Sciences Research Council grant “Physics of Fixational Eye Movements (PhysFEM)”, grant number EP/W023873/1.

## Notes

### Competing Interest Statement

The authors have declared no competing interest.

## Reference

[1] M. Rucci and M. Poletti, Control and functions of fixational eye movements, Annual review of vision science 1, 499 (2015).

[2] M. F. Land, Motion and vision: why animals move their eyes, Journal of Comparative Physiology A 185, 341 (1999).

[3] S. Martinez-Conde, S. L. Macknik, and D. H. Hubel, The role of fixational eye movements in visual perception, Nature reviews neuroscience 5, 229 (2004).

[4] R. G. Alexander and S. Martinez-Conde, Fixational eye movements, Eye movement research: An introduction to its scientific foundations and applications, 73 (2019).

[5] J. Jurin, A Reply to Mr. Robins’s Remarks on the Essay Upon Distinct and Indistinct Vision Published at the End of Dr. Smith’s Compleat System of Opticks: By James Jurin,… (W. Innys and R. Manby, 1739).

[6] R. Waring Darwin and E. Darwin, New experiments on the ocular spectra of light and colours. by robert waring darwin, md; communicated by erasmus darwin, mdfrs, Philosophical Transactions of the Royal Society of London Series I 76, 313 (1786).

[7] H. Von Helmholtz, Helmholtz’s treatise on physiological optics, Vol. 3 (Optical Society of America, 1925).

[8] O. Packer and D. R. Williams, Blurring by fixational eye movements, Vision research 32, 1931 (1992).

[9] J. Burge, Image-computable ideal observers for tasks with natural stimuli, Annual Review of Vision Science 6, 491 (2020).

[10] W. Bialek, Physical limits to sensation and perception, Annual review of biophysics and biophysical chemistry 16, 455 (1987).

[11] S. Hecht, S. Shlaer, and M. H. Pirenne, Energy, quanta, and vision, The Journal of general physiology 25, 819 (1942).

[12] D. A. Baylor, T. D. Lamb, and K.-W. Yau, Responses of retinal rods to single photons., The Journal of physiology 288, 613 (1979).

[13] A. Dey, A. J. Zele, B. Feigl, and P. Adhikari, Threshold vision under full-field stimulation: Revisiting the minimum number of quanta necessary to evoke a visual sensation, Vision Research 180, 1 (2021).

[14] H. B. Barlow, The size of ommatidia in apposition eyes, Journal of experimental Biology 29, 667 (1952).

[15] R. P. Feynman, R. B. Leighton, and M. Sands, The Feynman Lectures on Physics, Volume I (Addison-Wesley, Reading, Massachusetts, 1963).

[16] T. C. McLeish, Are there ergodic limits to evolution? ergodic exploration of genome space and convergence, Interface Focus 5, 20150041 (2015).

[17] E. N. Clarkson and R. Levi-Setti, Trilobite eyes and the optics of Descartes and Huygens, Nature 254, 663 (1975).

[18] E. Hering, Über die grenzen der sehschärfe, Ber. math.-phys. Cl. D. königl. Sächs. Gesell. Wiss. Leipzig 303, 16 (1899).

[19] M. Rucci, R. Iovin, M. Poletti, and F. Santini, Miniature eye movements enhance fine spatial detail, Nature 447, 852 (2007).

[20] K. Ratnam, N. Domdei, W. M. Harmening, and A. Roorda, Benefits of retinal image motion at the limits of spatial vision, Journal of vision 17, 30 (2017).

[21] J. Intoy and M. Rucci, Finely tuned eye movements enhance visual acuity, Nature communications 11, 795 (2020).

[22] J. Nachmias, Determiners of the drift of the eye during monocular fixation, Journal of the Optical Society of America 51, 761 (1961).

[23] R. M. Steinman, G. M. Haddad, A. A. Skavenski, and D. Wyman, Miniature eye movement: The pattern of saccades made by man during maintained fixation may be a refined but useless motor habit., Science 181, 810 (1973).

[24] E. Kowler, Eye movements: The past 25 years, Vision research 51, 1457 (2011).

[25] E. Ahissar and A. Arieli, Seeing via miniature eye movements: a dynamic hypothesis for vision, Frontiers in computational neuroscience 6, 89 (2012).

[26] M. Rucci, E. Ahissar, D. C. Burr, I. Kagan, M. Poletti, and J. D. Victor, The visual system does not operate like a camera, Journal of Vision 25, 2 (2025).

[27] R. Ditchburn and B. Ginsborg, Vision with a stabilized retinal image, Nature 170, 36 (1952).

[28] L. A. Riggs, F. Ratliff, J. C. Cornsweet, and T. N. Cornsweet, The disappearance of steadily fixated visual test objects, Journal of the Optical Society of America 43, 495 (1953).

[29] M. Rucci and J. D. Victor, The unsteady eye: an information-processing stage, not a bug, Trends in neurosciences 38, 195 (2015).

[30] L. E. Arend Jr, Spatial differential and integral operations in human vision: implications of stabilized retinal image fading., Psychological review 80, 374 (1973).

[31] H. L. Averill and F. W. Weymouth, Visual perception and the retinal mosaic. ii. the influence of eye-movements on the displacement threshold., Journal of Comparative Psychology 5, 147 (1925).

[32] W. Marshall and S. Talbot, Recent evidence for neural mechanisms in vision leading to a general theory of sensory acuity., (1942).

[33] R. M. Steinman and J. Z. Levinson, The role of eye movement in the detection of contrast and spatial detail, Eye movements and their role in visual and cognitive processes 4, 115 (1990).

[34] E. Ahissar and A. Arieli, Figuring space by time, Neuron 32, 185 (2001).

[35] U. Tulunay-Keesey and R. M. Jones, The effect of micromovements of the eye and exposure duration on contrast sensitivity, Vision research 16, 481 (1976).

[36] M. Gur, Seeing on the fly: Physiological and behavioral evidence show that space-to-space representation and processing enable fast and efficient performance by the visual system, Journal of Vision 24, 11 (2024).

[37] H. Barlow, Temporal and spatial summation in human vision at different background intensities, The Journal of physiology 141, 337 (1958).

[38] A. B. Watson, K. Boff, L. Kaufman, and J. Thomas, Handbook of perception and human performance (1986).

[39] M. Potters and W. Bialek, Statistical mechanics and visual signal processing, Journal de Physique I 4, 1755 (1994).

[40] D. A. Clark and J. E. Fitzgerald, Optimization in visual motion estimation, Annual review of vision science 10 (2024).

[41] W. Bialek, Biophysics: searching for principles (Princeton University Press, 2012).

[42] H. De Vries, The quantum character of light and its bearing upon threshold of vision, the differential sensitivity and visual acuity of the eye, Physica 10, 553 (1943).

[43] A. Rose, The relative sensitivities of television pickup tubes, photographic film, and the human eye, Proceedings of the IRE 30, 293 (1942).

[44] A. Rose, The sensitivity performance of the human eye on an absolute scale, Journal of the optical society of America 38, 196 (1948).

[45] See Supplemental Material at [URL will be inserted by publisher] for all derivations of key results, key measures of gaining information about external stimuli, and discussion of alternative forms of temporal filter.

[46] A. T. Rider, G. B. Henning, and A. Stockman, Light adaptation controls visual sensitivity by adjusting the speed and gain of the response to light, PloS one 14, e0220358 (2019).

[47] X. Kuang, M. Poletti, J. D. Victor, and M. Rucci, Temporal encoding of spatial information during active visual fixation, Current Biology 22, 510 (2012).

[48] J. Porter, A. Guirao, I. G. Cox, and D. R. Williams, Monochromatic aberrations of the human eye in a large population, Journal of the optical Society of America A 18, 1793 (2001).

[49] L. N. Thibos, X. Hong, A. Bradley, and X. Cheng, Statistical variation of aberration structure and image quality in a normal population of healthy eyes, Journal of the Optical Society of America A 19, 2329 (2002).

[50] J. G. Robson, Spatial and temporal contrast-sensitivity functions of the visual system, Journal of the optical society of America 56, 1141 (1966).

[51] F. W. Campbell and J. G. Robson, Application of fourier analysis to the visibility of gratings, The Journal of physiology 197, 551 (1968).

[52] R. L. De Valois, H. Morgan, and D. M. Snodderly, Psychophysical studies of monkey vision-iii. spatial luminance contrast sensitivity tests of macaque and human observers, Vision research 14, 75 (1974).

[53] D. H. Kelly, Motion and vision. ii. stabilized spatio-temporal threshold surface, Journal of the optical society of America 69, 1340 (1979).

[54] H. D. L. Dzn, Experiments on flicker and some calculations on an electrical analogue of the foveal systems, Physica 18, 935 (1952).

[55] A. Casile, J. D. Victor, and M. Rucci, Contrast sensitivity reveals an oculomotor strategy for temporally encoding space, Elife 8, e40924 (2019).

[56] Y. Burak, U. Rokni, M. Meister, and H. Sompolinsky, Bayesian model of dynamic image stabilization in the visual system, Proceedings of the National Academy of Sciences 107, 19525 (2010).

[57] M. Rucci and A. Casile, Fixational instability and natural image statistics: Implications for early visual representations, Network: Computation in Neural Systems 16, 121 (2005).

[58] M. Boi, M. Poletti, J. D. Victor, and M. Rucci, Consequences of the oculomotor cycle for the dynamics of perception, Current Biology 27, 1268 (2017).

[59] C. Enroth-Cugell and J. G. Robson, The contrast sensitivity of retinal ganglion cells of the cat, The Journal of physiology 187, 517 (1966).

[60] C. Enroth-Cugell, J. Robson, D. Schweitzer-Tong, and A. Watson, Spatio-temporal interactions in cat retinal ganglion cells showing linear spatial summation., The Journal of physiology 341, 279 (1983).

[61] L. J. Croner and E. Kaplan, Receptive fields of p and m ganglion cells across the primate retina, Vision research 35, 7 (1995).

[62] D. M. Dacey, Primate retina: cell types, circuits and color opponency, Progress in retinal and eye research 18, 737 (1999).

[63] L. Diller, O. S. Packer, J. Verweij, M. J. McMahon, D. R. Williams, and D. M. Dacey, L and m cone contributions to the midget and parasol ganglion cell receptive fields of macaque monkey retina, Journal of Neuroscience 24, 1079 (2004).

[64] J. Kulikowski and D. Tolhurst, Psychophysical evidence for sustained and transient detectors in human vision, The Journal of Physiology 232, 149 (1973).

[65] D. Lamming, Contrast sensitivity. In: Vision and visual dysfunction, ed. Cronly-Dillon, J., Vol.5 (London: Macmillan Press, 1991).

[66] W. Hart Jr, The temporal responsiveness of vision. In: Adlers Physiology of the Eye, Clinical Applicaiton, ed. Moses, R.J. and Hart, W.M. (St. Louis, The C.V. Mosby Company, 1987).

[67] D. Ruderman and W. Bialek, Statistics of natural images: Scaling in the woods, Advances in neural information processing systems 6 (1993).

[68] W. S. Geisler, Visual perception and the statistical properties of natural scenes, Annu. Rev. Psychol. 59, 167 (2008).

[69] E. G. Wu, N. Brackbill, C. Rhoades, A. Kling, A. R. Gogliettino, N. P. Shah, A. Sher, A. M. Litke, E. P. Simoncelli, and E. Chichilnisky, Fixational eye movements enhance the precision of visual information transmitted by the primate retina, Nature communications 15, 7964 (2024).

[70] F. Rieke, D. Warland, R. de Ruyter van Steveninck, and W. Bialek, Spikes: Exploring the Neural Code (MIT Press, Cambridge, MA, 1997).

[71] J. L. Reiniger, N. Domdei, F. G. Holz, and W. M. Harmening, Human gaze is systematically offset from the center of cone topography, Current Biology 31, 4188 (2021).

[72] J. A. Patrick, N. W. Roach, and P. V. McGraw, Motion-based super-resolution in the peripheral visual field, Journal of Vision 17, 15 (2017).

[73] J. L. Witten, V. Lukyanova, and W. M. Harmening, Sub-cone visual resolution by active, adaptive sampling in the human foveola, Elife 13, RP98648 (2024).

