## Supplemental Material for "Information, Movement and Adaptation in Human Vision"

### Information, Movement and Adaptation in Human Vision: Supplemental Material

[1]*School of Mathematics and Statistics, University of Glasgow, University Place, Glasgow G12 8QQ, United Kingdom*
[2]*Department of Psychology, University of Pennsylvania, Philadelphia, PA 19104-6241 USA*
[3]*Department of Experimental Psychology, University of Oxford, Oxford, OX2 6GG*
[4]*School of Mathematics, University of Leeds, Leeds, LS2 9JT, United Kingdom*
(Dated: Wednesday 3rd September, 2025)

#### I. GAINING INFORMATION ABOUT EXTERNAL STIMULUS FROM RETINAL RESPONSE

In section II A of the main paper we described a simple model for response of an array of photoreceptors. This is summarised in equation 1 of the main paper which gives the receptor response $y(\mathbf{r},t)$ from an external signal $x(\mathbf{r})$. In this section we discuss four different ways of thinking about how the response $y(\mathbf{r},t)$ is used to learn about the external signal $x(\mathbf{r})$ and develop the relevant calculations.

Our calculations are facilitated by taking the spatial Fourier transform of equation 1 of the main paper. This gives

$$\begin{aligned}\hat{y}_{\mathbf{k}}(t) &= \hat{F}_{\mathbf{k}}\hat{x}_{\mathbf{k}}\int_0^t \mathrm{d}t' A(t')K(t-t')e^{i\mathbf{k}\cdot\mathbf{R}(t')} + \hat{\eta}_{\mathbf{k}}(t)\\ &= \hat{F}_{\mathbf{k}}\hat{x}_{\mathbf{k}}b_{\mathbf{k}}(t) + \hat{\eta}_{\mathbf{k}}(t).\end{aligned} \tag{1}$$

where

$$b_{\mathbf{k}}(t) = \int_0^t \mathrm{d}t' A(t')K(t-t')e^{i\mathbf{k}\cdot\mathbf{R}(t')}. \tag{2}$$

To proceed, we take the measurements of $\hat{y}_{\mathbf{k}}(t)$ to occur at $n$ equally spaced times during observation time $t_{\mathrm{obs}}$, with time increment $\Delta t$:

$$t_i = \left(i - \frac{1}{2}\right)\Delta t, \qquad \Delta t = \frac{t_{\mathrm{obs}}}{N}, \qquad i = 1\ldots n, \tag{3}$$

so that

$$\hat{y}_{\mathbf{k}i} = \hat{F}_{\mathbf{k}}\hat{x}_{\mathbf{k}}b_{\mathbf{k}i} + \hat{\eta}_{\mathbf{k}i}, \tag{4}$$

where the $i$ subscript indicates that the function is evaluated at $t = t_i$. We will ultimately take the limits of $t_{\mathrm{obs}} \to \infty$ and $\Delta t \to 0$.

##### A. A naïve estimator for $\hat{x}_{\mathbf{k}}$

Eq. 4 is of form $\mathbf{y} = \hat{F}_{\mathbf{k}}\hat{x}_{\mathbf{k}}\mathbf{b} + \mathbf{n}$ where $\mathbf{y}$ is a vector whose $n$ components are observations $\hat{y}_{\mathbf{k}i}$ at times $t_i$, $\mathbf{b}$ is the vector whose components are $b_{\mathbf{k}i}$ and $\mathbf{n}$ is the vector of corresponding noise values $\hat{\eta}_{\mathbf{k}i}$. A naïve estimation of $\hat{x}_{\mathbf{k}}$ can be obtained by taking the scalar product $\mathbf{y}\cdot\mathbf{b}^*$ and rearranging for $\hat{x}_{\mathbf{k}}$. This gives:

$$\hat{x}_{\mathbf{k},\mathrm{est}} = \frac{\mathbf{y}\cdot\mathbf{b}^*}{\hat{F}_{\mathbf{k}}\left|\mathbf{b}\right|^2}. \tag{5}$$

*
†

Taking the limit of continuous observations $\Delta t \to 0$ means casting the scalar products as integrals, i.e. we estimate $\hat{x}_{\mathbf{k}}$ as:

$$\hat{x}_{\mathbf{k},\text{est}} = \frac{\int_0^{t_{\text{obs}}} \mathrm{d}t \hat{y}_{\mathbf{k}}(t) b_{\mathbf{k}}(t)^*}{\hat{F}_{\mathbf{k}} \int_0^{t_{\text{obs}}} \mathrm{d}t \left|b_{\mathbf{k}}(t)\right|^2}. \tag{6}$$

In what follows, for simplicity, we take the limit of long observation time, $t_{\text{obs}} \to \infty$.

This naïve estimator has the merit that it extracts only the component of measurements $\mathbf{y}$ in the direction of $\mathbf{b}$ (thus eliminating any parts of the noise that are incoherent with $\mathbf{b}$). It also prioritises measurements of $\hat{y}_{\mathbf{k}}(t)$ when the signal, via $b_{\mathbf{k}}(t)$, is large; conversely, where $b_{\mathbf{k}}(t)$ is zero or small, the measurement $\hat{y}_{\mathbf{k}}(t)$ is not used because the signal will be dominated by noise and so less informative.

We can consider the accuracy of this naïve estimator by evaluating the error, which is due to the noise:

$$\hat{x}_{\mathbf{k},\text{est}} - \hat{x}_{\mathbf{k}} = \frac{\mathbf{n} \cdot \mathbf{b}^*}{\hat{F}_{\mathbf{k}} \left|\mathbf{b}\right|^2} = \frac{\int_0^{\infty} \mathrm{d}t \hat{\eta}_{\mathbf{k}}(t) b_{\mathbf{k}}(t)^*}{\hat{F}_{\mathbf{k}} \int_0^{\infty} \mathrm{d}t \left|b_{\mathbf{k}}(t)\right|^2}. \tag{7}$$

Under the assumption of white noise, i.e. $\langle \hat{\eta}_{\mathbf{k}}(t) \hat{\eta}_{\mathbf{k}}(t') \rangle = N_{\mathbf{k}} \delta(t - t')$, we evaluate the mean square error to be

$$\langle \left|\hat{x}_{\mathbf{k},\text{est}} - \hat{x}_{\mathbf{k}}\right|^2 \rangle = \frac{N_{\mathbf{k}}}{\left|\hat{F}_{\mathbf{k}}\right|^2 \int_0^{\infty} \mathrm{d}t \left|b_{\mathbf{k}}(t)\right|^2}. \tag{8}$$

The form of this is easily understood. Each measurement $\hat{y}_{\mathbf{k}}(t)$ with non-zero $b_{\mathbf{k}}(t)$ carries some information about the external signal $\hat{x}_{\mathbf{k}}$, and so allows the signal to be estimated with greater accuracy. So, by accumulating measurements over time, the error in the estimate is reduced. The key quantity here is $\psi_{\mathbf{k}} = \int_0^T \mathrm{d}t \left|b_{\mathbf{k}}(t)\right|^2$ as defined in Eq. 2 of the main paper which indicates the combined effects of eye-movement, signal modulation, and adaptation on the accuracy of inference of the external signal. By making $\psi_{\mathbf{k}}$ as large as possible the error is minimised. It is clear that there is a benefit in making $b_{\mathbf{k}}(t)$ to be as large as possible for as long as possible.

##### B. Bayesian estimator for $\hat{x}_{\mathbf{k}}$

The analysis above was, however, on the basis of a naïve estimator of the external signal, which in general is not the optimal estimator. We can improve on the naïve estimator through a Bayesian analysis.

Assuming Gaussian white noise $\langle \hat{\eta}_{\mathbf{k}}(t) \hat{\eta}_{\mathbf{k}}(t') \rangle = N_{\mathbf{k}} \delta(t - t')$ gives $\langle \hat{\eta}_{\mathbf{k}i} \hat{\eta}_{\mathbf{k}j} \rangle = \frac{N_{\mathbf{k}}}{\Delta t} \delta_{ij}$ for the discretised measurements at times $t_i$. Hence the probability of obtaining a set of measurements $\hat{y}_{\mathbf{k}i}$ given external signal $\hat{x}_{\mathbf{k}}$ is obtained from the probability distribution of the noise:

$$\begin{aligned} P(\{\hat{y}_{\mathbf{k}i}\}|\hat{x}_{\mathbf{k}}) &= \frac{1}{\sqrt{(2\pi N_{\mathbf{k}}/\Delta t)^n}} \exp\left(-\frac{\Delta t}{2N_{\mathbf{k}}} \sum_i \left|\hat{\eta}_{\mathbf{k}i}\right|^2\right) \\ &= \frac{1}{\sqrt{(2\pi N_{\mathbf{k}}/\Delta t)^n}} \exp\left(-\frac{\Delta t}{2N_{\mathbf{k}}} \sum_i \left|\hat{y}_{\mathbf{k}i} - \hat{F}_{\mathbf{k}} \hat{x}_{\mathbf{k}} b_{\mathbf{k}i}\right|^2\right). \end{aligned} \tag{9}$$

This function is also the "likelihood" of $\hat{x}_{\mathbf{k}}$ given $\hat{y}_{\mathbf{k}i}$ and is maximised with respect to varying $\hat{x}_{\mathbf{k}}$ by the naïve estimator given above, i.e. the naïve estimator is a maximum likelihood estimation.

For simplicity, we assume a Gaussian prior distribution of input signal $\hat{x}_{\mathbf{k}}$ with zero mean and variance $\langle \left|\hat{x}_{\mathbf{k}}\right|^2 \rangle_0$, i.e.

$$P(\hat{x}_{\mathbf{k}}) = \frac{1}{\sqrt{(2\pi \langle \left|\hat{x}_{\mathbf{k}}\right|^2 \rangle_0)^n}} \exp\left(-\frac{\left|\hat{x}_{\mathbf{k}}\right|^2}{2\langle \left|\hat{x}_{\mathbf{k}}\right|^2 \rangle_0}\right). \tag{10}$$

We can calculate the probability of of obtaining a set of measurements $\hat{y}_{\mathbf{k}i}$ as

$$P(\{\hat{y}_{\mathbf{k}i}\}) = \int \mathrm{d}\hat{x}_{\mathbf{k}} P(\{\hat{y}_{\mathbf{k}i}\}|\hat{x}_{\mathbf{k}}) P(\hat{x}_{\mathbf{k}}). \tag{11}$$

Then following the set of observations $\hat{y}_{\mathbf{k}}(t)$ we obtain the posterior distribution for $\hat{x}_{\mathbf{k}}$ conditional on the observations made:

$$P(\hat{x}_{\mathbf{k}}|\{\hat{y}_{\mathbf{k}i}\}) = \frac{P(\{\hat{y}_{\mathbf{k}i}\}|\hat{x}_{\mathbf{k}}) P(\hat{x}_{\mathbf{k}})}{P(\{\hat{y}_{\mathbf{k}i}\})}. \tag{12}$$

This distribution is also Gaussian. Taking the limit $t_{\rm obs} \to \infty$ and $\Delta t \to 0$ the mean of the distribution is:

$$E\left(\hat{x}_{\mathbf{k}}|\hat{y}_{\mathbf{k}}(t)\right) = \frac{\int_0^\infty \mathrm{d}t \hat{y}_{\mathbf{k}}(t) b_{\mathbf{k}}(t)^*}{\hat{F}_{\mathbf{k}}\psi_{\mathbf{k}}} \frac{1}{1+\Upsilon_{\mathbf{k}}^{-1}} \tag{13}$$

and its variance is

$$\mathrm{Var}(\hat{x}_{\mathbf{k}}|\hat{y}_{\mathbf{k}}(t)) = \frac{\langle|\hat{x}_{\mathbf{k}}|^2\rangle_0}{1+\Upsilon_{\mathbf{k}}} \tag{14}$$

where $\Upsilon_{\mathbf{k}}$ is a measure of the effective signal to noise ratio:

$$\Upsilon_{\mathbf{k}} = \frac{\langle|\hat{x}_{\mathbf{k}}|^2\rangle_0 \left|\hat{F}_{\mathbf{k}}\right|^2}{N_{\mathbf{k}}}\psi_{\mathbf{k}}. \tag{15}$$

As before, $\psi_{\mathbf{k}} = \int_0^T \mathrm{d}t \left|b_{\mathbf{k}}(t)\right|^2$ as defined in Eq. 2 of the main paper.

As the signal to noise $\Upsilon_{\mathbf{k}}$ becomes large (that is, if the signal modulation and eye movements are such that $\psi_{\mathbf{k}}$ is large enough) then Eqs. 13 and 14 tend towards the naïve results Eq. 7 and 8 respectively. This represents the transition from low signal, where the best estimator and variance are dominated by the prior distribution, towards high signal, where the estimation and error are constrained primarily by the observations made. As with the naïve estimator, $\psi_{\mathbf{k}}$ is the key quantity indicating the combined effects of eye-movement, signal modulation, and adaptation on the accuracy of inference of the external signal.

In practice, a Gaussian prior may not be the most representative of true neural processing, for example see [1]. However, we anticipate that the transition from low signal (estimate dominated by the prior) towards high signal (estimation dominated by the likelihood function, related to the observations made) will remain for any prior.

##### C. Mutual information of stimulus and photoreceptor response

A robust quantifier of the information gained about the exterior scene $x(\mathbf{r})$ is the mutual information between the stimulus and the photoreceptor response. This mutual information is given by

$$I(y(\mathbf{r},t);x(\mathbf{r})) = \int \mathcal{D}[y] \int \mathcal{D}[x] P(x(\mathbf{r}),y(\mathbf{r},t)) \log_2\left[\frac{P(y(\mathbf{r},t)|x(\mathbf{r}))}{P(y(\mathbf{r},t))}\right], \tag{16}$$

where the integration is over all possible input and output signals. Here, $P(x(\mathbf{r}),y(\mathbf{r},t))$ is the joint probability distribution that the external signal is $x(\mathbf{r})$ and the photoreceptor response is $y(\mathbf{r},t)$; $P(y(\mathbf{r},t)|x(\mathbf{r}))$ is the probability of the photoreceptor response for a given external signal; and $P(y(\mathbf{r},t))$ is the overall probability of the photoreceptor response averaged over all possible external signals.

Calculation of the mutual information is also facilitated by taking the spatial Fourier transform of the signal (as before) and assuming a set of measurements at discrete time intervals, as in equation 4. Making the further assumption of Gaussian prior distribution of input signal $\hat{x}_{\mathbf{k}}$ with zero mean and variance $\langle|\hat{x}_{\mathbf{k}}|^2\rangle_0$, the mutual information becomes:

$$\begin{aligned} I(y;x) &= \sum_{\mathbf{k}} \int \mathrm{d}^n \hat{y}_{\mathbf{k}i} \int \mathrm{d}\hat{x}_{\mathbf{k}} P(\hat{x}_{\mathbf{k}}, \{\hat{y}_{\mathbf{k}i}\}) \log_2\left[\frac{P(\{\hat{y}_{\mathbf{k}i}\}|\hat{x}_{\mathbf{k}})}{P(\{\hat{y}_{\mathbf{k}i}\})}\right] \\ &= \sum_{\mathbf{k}} \frac{1}{2}\log_2\left(\frac{\mathrm{det}M_{y\mathbf{k}}}{\mathrm{det}M_{\eta\mathbf{k}}}\right), \end{aligned} \tag{17}$$

where the correlation matrices for the noise and the output are given by

$$\begin{aligned} (M_{\eta\mathbf{k}})_{ij} &= \langle\hat{\eta}_{\mathbf{k}i}\hat{\eta}_{\mathbf{k}j}\rangle \\ &= \frac{N_{\mathbf{k}}}{\Delta t}\delta_{ij} \end{aligned} \tag{18}$$

and

$$\begin{aligned} (M_{y\mathbf{k}})_{ij} &= \langle\hat{y}_{\mathbf{k}i}\hat{y}_{\mathbf{k}i}^*\rangle \\ &= |\hat{F}_{\mathbf{k}}|^2\langle|\hat{x}_{\mathbf{k}}|^2\rangle_0 b_{\mathbf{k}i}b_{\mathbf{k}j}^* + \frac{N_{\mathbf{k}}}{\Delta t}\delta_{ij}. \end{aligned} \tag{19}$$

The noise correlations are already fully diagonalised, making the determinant trivial to find, but $M_y$ is diagonal only in $\mathbf{k}$, not in time. To diagonalise it we notice that its eigenvectors are $b_{\mathbf{k}i}$ and $\{e_{\mathbf{k}\alpha,i}\}$, a set of $n-1$ vectors, with $\alpha = 1, \ldots, (n-1)$, perpendicular to $b_{\mathbf{k}i}$

$$(M_{y\mathbf{k}})_{ij}\, b_{\mathbf{k}j} = \left(|\hat{F}_{\mathbf{k}}|^2 \langle |\hat{x}_{\mathbf{k}}|^2 \rangle_0 |\mathbf{b}_{\mathbf{k}}|^2 + \frac{N_{\mathbf{k}}}{\Delta t}\right) b_{\mathbf{k}i}, \tag{20}$$

$$(M_{y\mathbf{k}})_{ij}\, e_{\mathbf{k}\alpha,j} = \frac{N_{\mathbf{k}}}{\Delta t} e_{\mathbf{k}\alpha,i} \qquad \forall \alpha. \tag{21}$$

The mutual information is then given by

$$I(y;x) = \sum_{\mathbf{k}} \frac{1}{2} \log_2 \left(1 + \Delta t \frac{|\hat{F}_{\mathbf{k}}|^2 \langle |\hat{x}_{\mathbf{k}}|^2 \rangle_0}{N_{\mathbf{k}}} |\mathbf{b}_{\mathbf{k}}|^2 \right), \tag{22}$$

which, in the limit $\Delta t \to 0$, becomes

$$I(y;x) = \sum_{\mathbf{k}} \frac{1}{2} \log \left(1 + \Upsilon_{\mathbf{k}}\right), \tag{23}$$

where $\Upsilon_{\mathbf{k}}$ is the effective signal to noise ratio defined above in equation 15, which includes the key quantity $\psi_{\mathbf{k}}$. We note that for small values of $\Upsilon_{\mathbf{k}} \ll 1$ the information grows linearly with increasing $\psi_{\mathbf{k}}$: this corresponds to the regime in the Bayesian estimator above where the prior dominates the estimate of $\hat{x}_{\mathbf{k}}$. For larger $\Upsilon_{\mathbf{k}} \gg 1$ the information grows logarithmically with increasing $\psi_{\mathbf{k}}$, which is the regime where the observations dominate the estimate of $\hat{x}_{\mathbf{k}}$.

##### D. Stimulus detection

Finally, we can draw some inspiration from theories of stimulus detection, i.e. whether a weak signal presented to the retina can be distinguished above the background noise. A simple model for stimulus detection was discussed by Watson [2] specifically in the context of the temporal response to an external signal. Watson considered the probability of detection of a stimulus over some observation time to be the probability that the signal is detected in at least one interval of time. He further used a simple relation between visual response and probability of detection in each time interval to arrive at a detection probability

$$p = 1 - \exp\left(-\alpha \sum_i |Y_i|^\beta\right) \tag{24}$$

where $Y_i$ is the (noise free) visual response in a given discrete time interval $i$, with the sum being over time intervals, $\alpha$ is a normalisation constant, and $\beta$ is a model parameter that can be chosen to reflect the steepness of the response curve. For present purposes, we make the specific choice $\beta = 2$. Although the particular functional form of Eq. 24 depends on some assumptions made about how probabilities of detection in given intervals combine, the general result that detection probability is an increasing function of $\sum_i |Y_i|^\beta$ seems not unreasonable, even if highly simplified. In our case we are considering a response at the retina which is a function of both time and retinal position, so we may extend the summation to include space and time. Taking the continuum limit, and assuming a long observation time

$$\begin{aligned} p &= 1 - \exp\left(-\alpha \int \mathrm{d}^2\mathbf{r} \int_0^\infty \mathrm{d}t\, |Y(\mathbf{r},t)|^2\right) \\ &= 1 - \exp\left(-\frac{\alpha}{4\pi^2} \int \mathrm{d}^2\mathbf{k} \int_0^\infty \mathrm{d}t \left|\hat{Y}_{\mathbf{k}}(t)\right|^2\right) \end{aligned} \tag{25}$$

where, as before, we take the spatial Fourier transform of the signal. Taking $\hat{Y}_{\mathbf{k}}(t) = A\hat{F}_{\mathbf{k}}\hat{x}_{\mathbf{k}} b_{\mathbf{k}}(t)$ as the noise free signal, with $A$ being an overall multiplier of stimulus intensity that can be varied to find the detection threshold,

$$\begin{aligned} p &= 1 - \exp\left(-A^2 \frac{\alpha}{4\pi^2} \int \mathrm{d}^2\mathbf{k} \left|\hat{F}_{\mathbf{k}}\right|^2 |\hat{x}_{\mathbf{k}}|^2 \int_0^\infty \mathrm{d}t\, |b_{\mathbf{k}}(t)|^2\right) \\ &= 1 - \exp\left(-A^2 \frac{\alpha}{4\pi^2} \int \mathrm{d}^2\mathbf{k} \left|\hat{F}_{\mathbf{k}}\right|^2 |\hat{x}_{\mathbf{k}}|^2 \psi_{\mathbf{k}}\right). \end{aligned} \tag{26}$$

Here we see that, within the approximation of the Watson signal detection model, and further with the specific choice $\beta = 2$, the probability of signal detection increases with increasing $\psi_{\mathbf{k}}$, lending further support towards the maximisation of this quantity. Further, this specific model indicates how $\psi_{\mathbf{k}}$ might be combined with signal $\hat{x}_{\mathbf{k}}$ to optimise that detection.

In Eq. 21 in the main text we replace $\psi_{\mathbf{k}}$ with its average value $\langle\psi_{\mathbf{k}}\rangle$, where the average is taken over eye motion paths. We now discuss this averaging and derive a convenient expression for $\chi_{\mathbf{k}}$.

#### II. AVERAGING OVER EYE MOTION TRAJECTORIES

In the previous section we presented several approaches to calculating the information gained about an external stimulus from the retinal response in the presence of eye movement. In all cases, the key quantity to calculate was $\psi_{\mathbf{k}} = \int_0^T \mathrm{d}t\, |b_{\mathbf{k}}(t)|^2$. The value of $\psi_{\mathbf{k}}$ depends on the specific path $\mathbf{R}(t')$ of the eye motion, i.e. it takes a different value for different paths. In this paper we focus on its average $\langle\psi_{\mathbf{k}}\rangle$ across multiple paths, reasoning that forms of eye motion that maximise the average are likely to be beneficial.

It is possible in principle to probe the consequences of focussing only on the average value of $\psi_{\mathbf{k}}$. For example, when calculating the mutual information, we may be interested in averaging this quantity over the eye trajectories, i.e. in calculating:

$$\langle I(y;x)\rangle_{\mathbf{R}} = \Big\langle \sum_{\mathbf{k}} \frac{1}{2} \log\left(1 + \Upsilon_{\mathbf{k}}\right) \Big\rangle_{\mathbf{R}}, \tag{27}$$

where $\langle \bullet \rangle_{\mathbf{R}}$ denotes an average over paths. The contribution from these paths is entirely within the quantity $\Upsilon_{\mathbf{k}}$. To handle the expectation of a logarithm we use that for a sufficiently differentiable function $f$

$$\langle f(X)\rangle \approx f(\mu_X) + \frac{1}{2} f''(\mu_X)\sigma_X, \tag{28}$$

where $\mu_X$ and $\sigma_X$ are the mean and variance respectively of the random variable $X$. Hence:

$$\langle I(y;x)\rangle_{\mathbf{R}} \approx \sum_{\mathbf{k}} \frac{1}{2}\left[\log_2\left(1 + \langle\Upsilon_{\mathbf{k}}\rangle_{\mathbf{R}}\right) - \frac{1}{2}\frac{\langle(\Upsilon_{\mathbf{k}} - \langle\Upsilon_{\mathbf{k}}\rangle_{\mathbf{R}})^2\rangle_{\mathbf{R}}}{\left(1 + \langle\Upsilon_{\mathbf{k}}\rangle_{\mathbf{R}}\right)^2}\right]. \tag{29}$$

In this paper we are effectively considering only the first term, which depends on $\langle\Upsilon_{\mathbf{k}}\rangle_{\mathbf{R}} = \frac{\langle|\hat{x}_{\mathbf{k}}|^2\rangle_0 |\hat{F}_{\mathbf{k}}|^2}{N_{\mathbf{k}}}\langle\psi_{\mathbf{k}}\rangle$. This gives an upper bound on the mutual information provided by Jensen's inequality. It is possible to quantify the error in this approximation via the variance of $\Upsilon_{\mathbf{k}}$, in the second term of Eq. (29).

##### A. Calculating $\langle\psi_{\mathbf{k}}\rangle$

We wish to calculate the average over eye trajectories of $\psi_{\mathbf{k}} = \int_0^T \mathrm{d}t\, |b_{\mathbf{k}}(t)|^2$, which is

$$\langle\psi_{\mathbf{k}}\rangle = \left\langle \int_0^\infty \mathrm{d}t \int_0^t \mathrm{d}t' \int_0^t \mathrm{d}t'' A(t')A(t'')K(t-t')K(t-t'')e^{i\mathbf{k}\cdot(\mathbf{R}(t')-\mathbf{R}(t''))} \right\rangle_{\mathbf{R}}. \tag{30}$$

Now, defining explicitly that both $A(t) = 0$ and $K(t) = 0$ for $t < 0$ allows us to extend the integration range of all three integrals to $(-\infty, \infty)$. Further, assuming that the dynamics of $\mathbf{R}(t)$ is statistically stationary (i.e. statistically independent of absolute time), allows us to define the path 'structure factor' function $S_{\mathbf{k}}(\tau) = \langle e^{i\mathbf{k}\cdot(\mathbf{R}(t+\tau)-\mathbf{R}(t))}\rangle$, with the average being over all trajectories, over all times $t$. This gives:

$$\langle\psi_{\mathbf{k}}\rangle = \int_{-\infty}^\infty \mathrm{d}t \int_{-\infty}^\infty \mathrm{d}t' \int_{-\infty}^\infty \mathrm{d}t'' A(t')A(t'')K(t-t')K(t-t'')S_{\mathbf{k}}(t'-t''). \tag{31}$$

Now making the substitutions $t = t' + u$ and $t' = t'' + \tau$ gives:

$$\begin{aligned}\langle\psi_{\mathbf{k}}\rangle &= \int_{-\infty}^\infty \mathrm{d}u \int_{-\infty}^\infty \mathrm{d}\tau \int_{-\infty}^\infty \mathrm{d}t'' A(t''+\tau)A(t'')K(u)K(u+\tau)S_{\mathbf{k}}(\tau) \\ &= \int_{-\infty}^\infty \mathrm{d}\tau C_{AA}(\tau)C_{KK}(\tau)S_{\mathbf{k}}(\tau).\end{aligned} \tag{32}$$

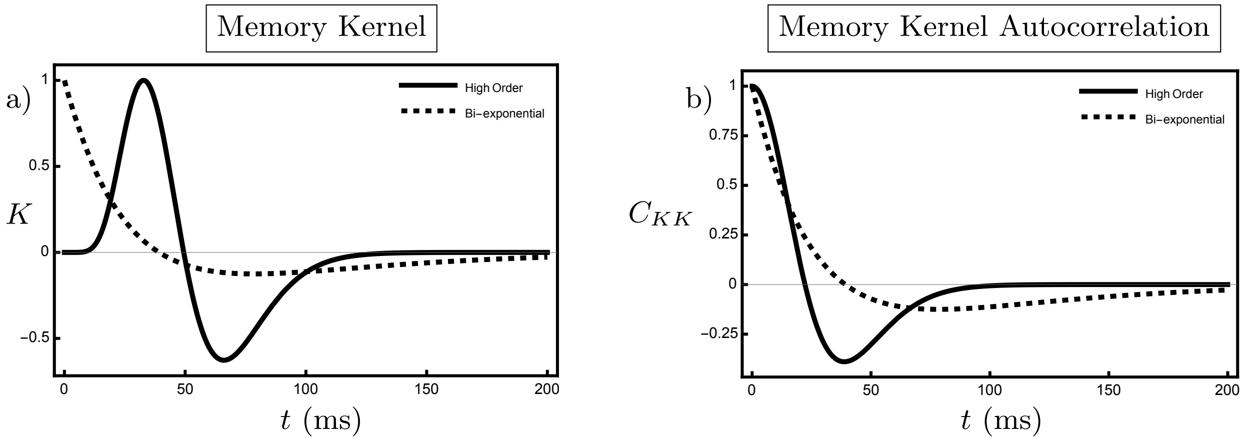


FIG. 1. The comparison between different forms of memory kernel. a) A high-order memory kernel, resulting from (35) with $n_1 = 9$ and $n_2 = 10$, is shown (solid) along with the bi-exponential kernel given by (34) (dashed). For the former $\sigma = 1.15$ and $t_a = 5.7$ ms, while for the bi-exponential kernel $\sigma = \sqrt{2}$ and $t_a = 40$ ms, as used in Figure 3c) and Figures 5 and 6. b) The corresponding autocorrelation functions with the same parameter choices as in a). All four functions have been normalised by their maximum value.

We note that all of $C_{AA}(\tau)$, $C_{KK}(\tau)$ and $S_{\mathbf{k}}(\tau)$ are even functions of $\tau$ allowing us to halve the integration domain. For convenience, we further normalise $\langle\psi_{\mathbf{k}}\rangle$ by the adaptation time $t_a$ of the memory kernel, giving (finally)

$$\chi_{\mathbf{k}} = \frac{\langle\psi_{\mathbf{k}}\rangle}{t_a} = \frac{2}{t_a}\int_0^\infty \mathrm{d}\tau C_{AA}(\tau)C_{KK}(\tau)S_{\mathbf{k}}(\tau), \tag{33}$$

as given in the main paper.

#### III. HIGHER-ORDER TEMPORAL FILTER

In the main text we have incorporated adaptation via a bi-exponential linear temporal response

$$K(\tilde{t}) = \frac{1}{t_a}\left[\sigma \mathrm{e}^{-\sigma\tilde{t}} - \frac{1}{\sigma}\mathrm{e}^{-\tilde{t}/\sigma}\right]. \tag{34}$$

Generalised forms of this response function have been proposed [2], on the grounds that they provide good agreement with observed human contrast sensitivity functions [3]. These memory kernels have the form

$$K(\tilde{t}) = \frac{1}{t_a}\left[\frac{1}{(n_1-1)!}\left(\sigma\tilde{t}\right)^{n_1-1}\sigma \mathrm{e}^{-\sigma\tilde{t}} - \frac{1}{(n_2-1)!}\left(\frac{\tilde{t}}{\sigma}\right)^{n_2-1}\frac{1}{\sigma}\mathrm{e}^{-\tilde{t}/\sigma}\right], \tag{35}$$

where the excitatory and inhibitory components result from a cascade of $n_1$ and $n_2$ identical low-pass filters respectively. We argue here that these generalisations do not qualitatively change the results presented in the main text, achieved with the more tractable bi-exponential kernel given in (34), which corresponds to the case $n_1 = n_2 = 1$. The key point is that even when the memory kernel changes considerably, its autocorrelation is altered much less, and it is the latter which enters the measure of acquired visual information, $\chi_{\mathbf{k}}$, via Eq. 3 of the main text.

We illustrate this point with the particular form of (35) used in [2], that is with $n_1 = 9$ and $n_2 = 10$. This gives a memory kernel shown by the solid line in Figure 1a), which unlike the dashed line representing the bi-exponential memory kernel, vanishes at the origin and achieves its maximum at a non-zero temporal delay. However these differences disappear in the corresponding autocorrelation functions, shown in Figure 1b), both of which are maximal at the origin. Since it is the autocorrelation function, $C_{KK}$ which is relevant for the determining $\chi_{\mathbf{k}}$, this suggests it is sufficient to use a bi-exponential memory kernel, provided the adaptation time is suitably scaled to account for the additional filters contained in (35). Note that in the notation of [2], $\tau$ is equivalent to our $t_s$, while $\kappa$ is equivalent to $\sigma^2$. Hence, $\tau = 4.3$ ms and $\kappa = 1.33$, as used to [2] to match the data of [3], corresponds to $\sigma = \sqrt{1.33}$ and $t_a = 4.3\sqrt{1.33}$, these being the values used for the plotting of (35) in Figure 1.

#### IV. IMPERFECT ADAPTATION

To include the possibility of imperfect adaptation we modify the memory kernel to be

$$K(t) = \frac{1}{\tau_s}\mathrm{e}^{-t/\tau_s} - \frac{\zeta}{\tau_l}\mathrm{e}^{-t/\tau_l}, \tag{36}$$

such that its integral tends to $1-\zeta$. The corresponding auto-correlation function is given by

$$C_{KK}(\tilde{\tau}) = \frac{1}{2t_a}\left[\sigma\mathrm{e}^{-\sigma\tilde{\tau}} - \frac{2\sigma}{\sigma^2+1}\zeta\left(\mathrm{e}^{-\sigma\tilde{\tau}} + \mathrm{e}^{-\tilde{\tau}/\sigma}\right) + \frac{1}{\sigma}\zeta^2\mathrm{e}^{-\tilde{\tau}/\sigma}\right], \tag{37}$$

such that, for stimuli of constant amplitude, the gained information is given by

$$\begin{aligned}\chi_{\mathbf{k}} &= \int_0^{\tilde{T}} (\tilde{T}-\tilde{\tau})\left[\sigma\mathrm{e}^{-\sigma\tilde{\tau}} - \frac{2\sigma}{\sigma^2+1}\zeta\left(\mathrm{e}^{-\sigma\tilde{\tau}} + \mathrm{e}^{-\tilde{\tau}/\sigma}\right) + \frac{1}{\sigma}\zeta^2\mathrm{e}^{-\tilde{\tau}/\sigma}\right]\mathrm{e}^{-\Gamma\tilde{\tau}}\mathrm{d}\tilde{\tau} \\ &= \frac{1}{2\left(\sigma^2+1\right)}\left\{\frac{\sigma\left(1-2\zeta+\sigma^2\right)}{\Gamma+\sigma}\left[\tilde{T} - \frac{1}{\Gamma+\sigma}\left(1-\mathrm{e}^{-(\Gamma+\sigma)\tilde{T}}\right)\right] + \frac{\zeta\left(\zeta+(\zeta-2)\sigma^2\right)}{1+\Gamma\sigma}\left[\tilde{T} - \frac{\sigma}{1+\Gamma\sigma}\left(1-\mathrm{e}^{-\left(\Gamma+\frac{1}{\sigma}\right)\tilde{T}}\right)\right]\right\}.\end{aligned} \tag{38}$$

This informatic gain is plotted in Figure 2a)-e), with Figure 2f) showing the corresponding asymptotic rate of information gain, namely the coefficient of $\tilde{T}$ in (38):

$$\frac{1}{2}\left(\frac{\sigma}{\Gamma+\sigma} + \frac{\zeta^2}{1+\Gamma\sigma}\right) - \frac{\zeta\sigma}{\sigma^2+1}\left(\frac{1}{\Gamma+\sigma} + \frac{\sigma}{1+\Gamma\sigma}\right). \tag{39}$$

Figure 2a) shows the case of perfect adaptation, also shown in Figure 2a) of the main text, illustrating the $1/\Gamma$ and $\Gamma$ power laws for small and large $\Gamma$ respectively, as discussed in Section III A of the main text. While the former remains in the subsequent panels, the low $\Gamma$ power law is lost, replaced by a constant rate of information gain. Indeed, expanding (39) around $\Gamma = 0$ gives

$$\frac{1}{2}\left(\zeta-1\right)^2 + \left(\frac{\zeta(\sigma^4+1)}{\sigma(\sigma^2+1)} - \frac{\zeta^2\sigma^2+1}{2\sigma}\right)\Gamma. \tag{40}$$

The constant term represents the steady leak of information through the adaptive filter due to imperfect adaptation. This effect of imperfect adaptation at low $\Gamma$ mirrors that of stimulus modulation seen in Section III B of the main text. In both cases perfect adaptation is avoided such that a finite rate of information gain is achieved asymptotically, either due to the adaptation being imperfect, or because temporal modulation circumvents adaptation, as discussed in Section III B of the main text.

Without the same need to overcome adaptation, it is not clear that retinal motion will still have a benefit, indeed one is not discernable in Figure 2b)-d). The linear coefficient in (40) allows us to quantify when an informatic benefit will arise due to retinal motion. Requiring this coefficient to be positive, we find retinal motion will remain beneficial despite imperfect adaptation provided

$$\sigma > \sqrt{\frac{\zeta^2+1+(1-\zeta)\sqrt{1+10\zeta+\zeta^2}}{2\zeta(2-\zeta)}}. \tag{41}$$

#### V. SINUSOIDAL MODULATION

We take a sinusoidally modulated stimulus to have amplitude $A(\tilde{t}) = \cos\left(\tilde{\omega}\tilde{t}\right)$ over a presentation time $\tilde{T}$, where, as in the main text, the tildes denotes non-dimensionalisation by the adaptation time $t_a$. The autocorrelation function of the modulation is then given by

$$C_{AA}(\tilde{\tau}) = \begin{cases}\frac{1}{2}t_a\left(\tilde{T}-|\tilde{\tau}|\right)\left[\cos\left(\tilde{\omega}|\tilde{\tau}|\right) + \cos\left(\tilde{\omega}\tilde{T}\right)\mathrm{sinc}\left(\tilde{\omega}\left(\tilde{T}-|\tilde{\tau}|\right)\right)\right] & 0 \le |\tilde{\tau}| \le \tilde{T} \\ 0 & \text{otherwise}\end{cases}. \tag{42}$$

This is illustrated in Figure 3 for the case $\tilde{\omega}\tilde{T} = 4\pi$, along with the amplitude autocorrelation for an unmodulated signal for comparison. As can be seen from (42), the amplitude autocorrelation contains oscillations at the same frequency as the signal modulation, within an envelope of half the value of the unmodulated autocorrelation, that is $(\tilde{T} - \tilde{\tau})/2$.

The gained information is then found via the analagous equation to Eq 10 of the main text, that is

$$\begin{aligned}
\chi_{\mathbf{k}} &= \frac{\sigma^2 - 1}{\sigma^2 + 1} \int_0^{\tilde{T}} \frac{1}{2}\left(\tilde{T} - \tilde{\tau}\right)\left[\cos\left(\tilde{\omega}\tilde{\tau}\right) + \cos\left(\tilde{\omega}\tilde{T}\right)\operatorname{sinc}\left(\tilde{\omega}\left(\tilde{T} - \tilde{\tau}\right)\right)\right]\left[\sigma \mathrm{e}^{-\sigma\tilde{\tau}} - \frac{1}{\sigma}\mathrm{e}^{-\tilde{\tau}/\sigma}\right]\mathrm{e}^{-\Gamma\tilde{\tau}}\mathrm{d}\tilde{\tau} \\
&= \frac{\sigma^2 - 1}{2(\sigma^2 + 1)}\left[\left(\frac{\sigma\left(\Gamma + \sigma\right)}{A_+} - \frac{1 + \Gamma\sigma}{B_+}\right)\tilde{T} + \sigma\left(\frac{B_-}{B_+^2} - \frac{A_-}{A_+^2}\right) + \sigma\left(\frac{1}{B_+} - \frac{1}{A_+}\right)\cos^2\left(\tilde{\omega}\tilde{T}\right)\right. \\
&+ \frac{1}{\tilde{\omega}}\left(\frac{\sigma\left(\Gamma + \sigma\right)}{A_+} - \frac{1 + \Gamma\sigma}{B_+}\right)\cos\left(\tilde{\omega}\tilde{T}\right)\sin\left(\tilde{\omega}\tilde{T}\right) \\
&+ \left(\frac{2\sigma\left(\Gamma + \sigma\right)^2 \mathrm{e}^{-(\Gamma + \sigma)\tilde{T}}}{A_+^2} - \frac{2\sigma\left(1 + \Gamma\sigma\right)^2 \mathrm{e}^{-\left(\Gamma + \frac{1}{\sigma}\right)\tilde{T}}}{B_+^2}\right)\cos\left(\tilde{\omega}\tilde{T}\right) \\
&\left.\sigma\tilde{\omega}\left(\frac{2\sigma\left(1 + \Gamma\sigma\right)\mathrm{e}^{-\left(\Gamma + \frac{1}{\sigma}\right)\tilde{T}}}{B_+^2} - \frac{2\left(\Gamma + \sigma\right)\mathrm{e}^{-(\Gamma + \sigma)\tilde{T}}}{A_+^2}\right)\sin\left(\tilde{\omega}\tilde{T}\right)\right],
\end{aligned} \tag{43}$$

where we have defined

$$\begin{aligned}
A_\pm &= \left(\Gamma + \sigma\right)^2 \pm \tilde{\omega}^2, \\
B_\pm &= \left(1 + \Gamma\sigma\right)^2 \pm \sigma^2\tilde{\omega}^2.
\end{aligned} \tag{44}$$

This expression for the visual information is shown in Figure 4 of the main text for a variety of regimes of oscillation frequencies and presentation times.


[1] E. G. Wu, N. Brackbill, C. Rhoades, A. Kling, A. R. Gogliettino, N. P. Shah, A. Sher, A. M. Litke, E. P. Simoncelli, and E. Chichilnisky, Fixational eye movements enhance the precision of visual information transmitted by the primate retina, Nature communications **15**, 7964 (2024).

[2] A. B. Watson, K. Boff, L. Kaufman, and J. Thomas, Handbook of perception and human performance (1986).

[3] H. D. L. Dzn, Experiments on flicker and some calculations on an electrical analogue of the foveal systems, Physica **18**, 935 (1952).

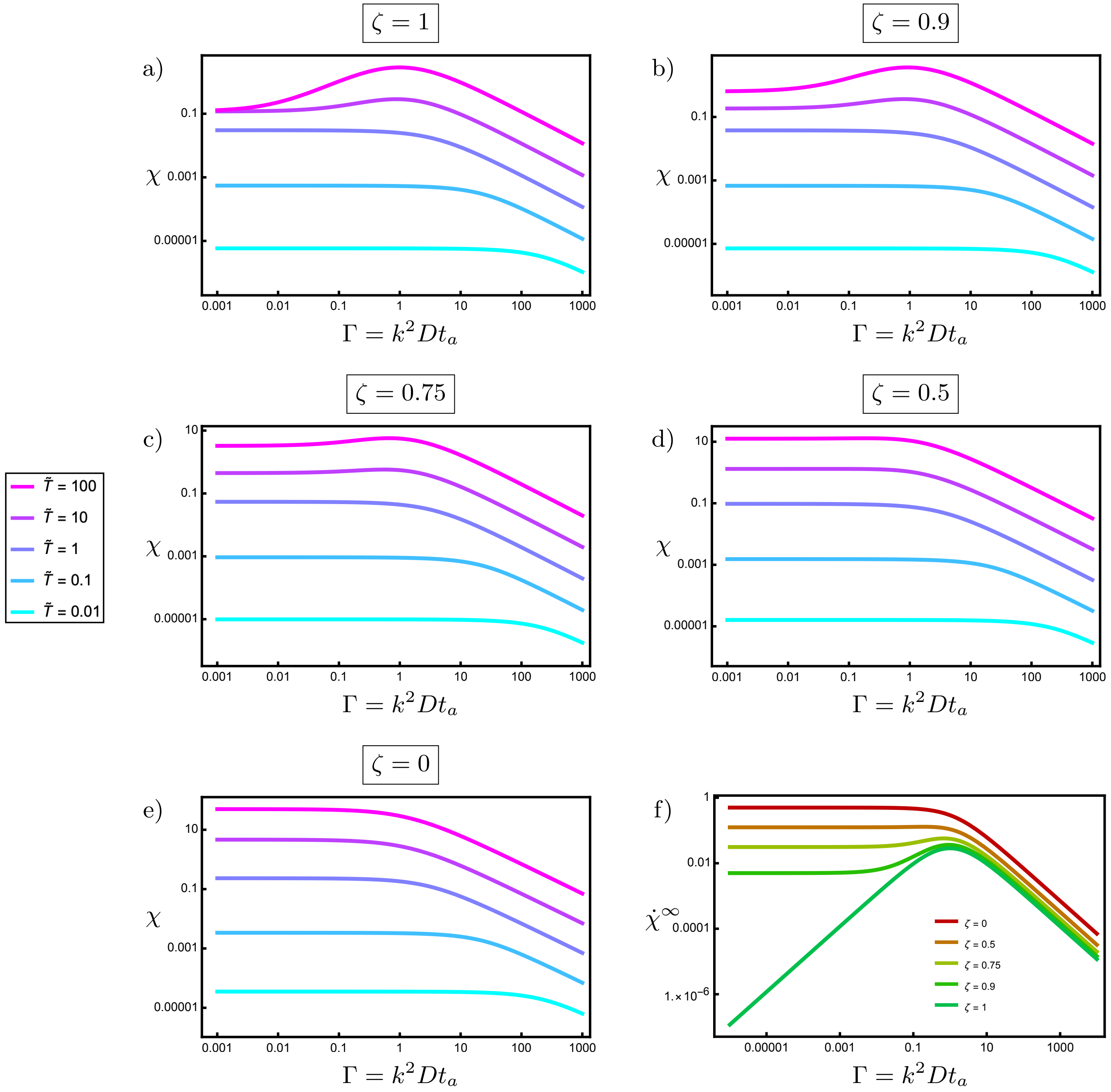


FIG. 2. The retinal information gained from a constant amplitude signal with diffusive ocular drift. The three components, namely ocular drift, stimulus and adaptation are captured through dimensionless parameters. The nature of the drift is described by $\Gamma$, shown on the horizontal axis, that is how far the retina diffuses in the adaptation time, measured in the wavelength of the signal. The form of the memory kernel is captured by a shape parameter $\sigma = \sqrt{t_l/t_s}$ and the stimulus is presented for a time $T = \tilde{T}t_a$. a) to e) The gained retinal information, as measured by $\chi_{\mathbf{k}}$, given in (38), with $\sigma = \sqrt{2}$ and $\tilde{T}$ varying from 0.01 (cyan) to 100 (magenta) in powers of 10. f) The corresponding asymptotic rates of information gain, $\dot{\chi}^{\infty}$, as provided in (39).

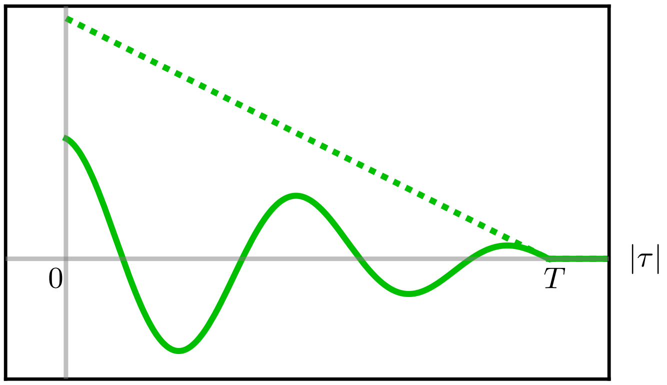


FIG. 3. The amplitude autocorrelation function for a stimulus modulated at frequency $\omega = \tilde{\omega}/t_a$ and presented for a time $T = \tilde{T} t_a$, with $t_a$ the adaptation time. The solid curve shows the amplitude autocorrelation when $\tilde{\omega}\tilde{T} = 4\pi$, while the dashed curve illustrates the case of an unmodulated signal, also shown in Figure 1e).
